# Tissue-Specific Iron Levels Modulate Lipid Peroxidation and the FLASH Radiotherapy Effect

**DOI:** 10.1101/2025.05.14.653978

**Authors:** Nuria Vilaplana-Lopera, Jiyoung Kim, Gilyeong Nam, Iain D. C. Tullis, Salome Paillas, Jia-Ling Ruan, Pei Ju Lee, Yanyan Jiang, Sohee Park, Tianxu Hou, Ayesha Nasir, Eve Charlesworth, Ellie Walker, Ammar Abu-Halawa, Mark A. Hill, Changhoon Choi, Ik Jae Lee, Youngtae Jeong, Samira Lakhal-Littleton, Chee Kin Then, Shing-Chuan Shen, Amato J. Giaccia, Kristoffer Petersson, Eui Jung Moon

**Affiliations:** Department of Oncology, University of Oxford, Oxford, United Kingdom; Department of Radiation Oncology, Severance Hospital, Yonsei University Medical School, Seoul, Republic of Korea; Graduate Institute of Medical Sciences, College of Medicine, Taipei Medical University, Taipei, Taiwan; Department of Radiation Oncology, Samsung Medical Center, Seoul, Republic of Korea; Department of New Biology, Daegu Gyeongbuk Institute of Science and Technology (DGIST), Daegu, Republic of Korea; Department of Physiology, Anatomy & Genetics, University of Oxford, Oxford, United Kingdom; Department of Radiation Oncology, Shuang Ho Hospital, Taipei Medical University, New Taipei City, Taiwan; Graduate Institute of Clinical Medicine, College of Medicine, Taipei Medical University, Taipei, Taiwan

## Abstract

Iron is vital to living cells, playing a key role in cellular respiration, DNA synthesis, and various metabolic functions. Importantly, cancer cells have a higher dependency on iron compared to normal cells to support their rapid growth and survival. Due to this fact, tumors are more vulnerable to ferroptosis, an iron-dependent form of regulated cell death. Radiation therapy (RT), a standard treatment for many cancer patients, is known to induce ferroptosis. Ultra-high dose rate FLASH RT offers an improved therapeutic window by minimizing damage to normal tissues while preserving tumor control. However, the precise biological mechanisms behind the protective effects of FLASH RT on normal tissues remain unclear. In this study, we propose that variations in lipid peroxidation and ferroptosis, driven by intrinsic differences in iron levels between normal and cancerous tissues, contribute to this effect. Our findings show that FLASH RT increases lipid peroxidation and induces ferroptosis in tumor cells but does not significantly elevate lipid peroxidation and ferroptosis in normal tissues compared to conventional RT. To determine whether raising iron levels in normal tissues could abrogate the protective effects of FLASH, mice were fed a high-iron diet before RT. A high-iron diet before and after RT reversed the protective effect of FLASH, resulting in increased intestinal damage and lipid peroxidation. This suggests that baseline iron levels and iron-driven lipid peroxidation are critical factors in mediating the protective outcomes of FLASH RT. Overall, our study sheds light on the role of iron in modulating RT responses and provides new mechanistic insights into how FLASH RT influences normal and cancerous tissues.

## Introduction

Iron is a vital element that plays a fundamental role in maintaining essential cellular functions such as oxygen transport, energy production, hematopoiesis, and DNA replication (1). Its biological importance becomes particularly essential in cancer, where tumor cells exhibit a strong dependence on iron for survival. Therefore, elevated levels of iron and ferritin, the primary intracellular iron storage protein, are frequently observed in the tissues and serum of cancer patients (2–5). This iron dependency opens a critical therapeutic window through ferroptosis, an iron-dependent form of regulated cell death characterized by excessive lipid peroxidation (6).

Radiation therapy (RT), a standard care for many cancer patients to achieve local control and to treat metastatic burden, has been shown to promote ferroptosis by inducing lipid peroxidation, thereby enhancing tumor cell death (7). Furthermore, combining RT with ferroptosis inducers such as RSL3, erastin, and sulfasalazine, has been shown to increase tumor radiosensitivity and overcome radioresistance (7–9). While RT works by damaging tumor DNA either directly or indirectly via the generation of reactive oxygen species (ROS), ferroptosis operates independently of DNA damage pathways, making it a promising approach for developing therapeutic strategies to increase RT response (10).

FLASH RT uses ultra-high dose rates to increase the therapeutic window of RT (11). While conventional RT is delivered at dose rates ranging from 0.01 to 0.4 Gy/s, FLASH RT, which is delivered at rates greater than 40 Gy/s, results in less normal tissue toxicity while maintaining tumor control rates comparable to conventional RT (12, 13). Normal tissue sparing after FLASH RT is known as the FLASH effect and has been attributed in part to the interplay between oxygen depletion and ROS production (11, 14–16). Although studies have demonstrated that FLASH RT reduces tissue oxygen concentrations both in normal tissues and tumors, oxygen depletion may not be the only mechanism accounting for normal tissue sparing, as the observed reduction in oxygen could be considered insufficient to induce tissue protection (17, 18). It has also been suggested that radical-radical recombination as well as immune modulation are potential mechanisms of FLASH RT (16, 19–21). Recently, the contribution of lipids and lipid peroxidation was proposed as the key link between FLASH and ferroptosis, an iron-dependent regulated cell death pathway (22–24). However, the biological mechanisms by which FLASH RT alters lipid peroxidation and potentially ferroptosis remain unclear. Therefore, this study investigated how lipid peroxidation and ferroptosis are differentially modulated after FLASH RT in tumor and normal tissues and whether varying iron levels in these tissues plays a role in the FLASH effect.

## Materials and Methods

### Cell lines and cell culture

A549 and MDA-MB-231 were purchased from ATCC and maintained at 37 °C in a humidified incubator with 5% CO_2_ in Dulbecco’s Modified Eagle Medium supplemented with 10% fetal bovine serum (F7524, Sigma Aldrich) and 1% antibiotic/antimycotic (15240-062, Gibco). Cells were routinely tested for mycoplasma.

### siRNA transfection

ONTARGETplus SMARTpool siRNA targeting TFRC (Horizon Discovery) was transfected into A549 and MDA-MB-231 cells according to the manufacturer’s instructions. Briefly, 2 × 10 A549 and MDA-MB-231 cells were seeded in 6-well plates. After 24 hours of incubation, the cells were washed and cultured in 1.6 mL of antibiotic-free DMEM supplemented with 10% FBS. A siRNA solution was prepared by diluting 10 µL of 20 µM TFRC-targeting siRNA in 190 µL of Opti-MEM (31985070, GIBCO), while 7.5 µL of RNAiMax (13778030, Thermo Fisher Scientific) was diluted in 192.5 µL of Opti-MEM. The two diluted components were then combined and incubated at room temperature for 15 minutes before being added to the cells. Cells were incubated overnight before being seeded for further experiments.

### Real Time quantitative reverse transcription PCR (qRT-PCR)

Total mRNA was extracted using TRIzolTM Reagent (15596026, Invitrogen) following manufacturer’s instructions. One μg of mRNA was reverse transcribed into cDNA using an iScript cDNA synthesis kit (1708890, BioRad). Quantitative real-time RT-PCR was performed with an iTaq^TM^ Universal SYBR Green Supermix (1725124, BioRad) using the StepOnePlus Real-Time PCR system (Applied Biosystems). Expression level of TFRC mRNA (Forward: GTTGAATTGAACCTGGAC, Reverse: AAGTAGCACGGAAGAAGT) was determined using the ΔΔCT method and was normalized to 18S expression (Forward: GAGGATGAGGTGGAACGTGT, Reverse: AGAAGTGACGCAGCCCTCTA) in the same sample.

### Intracellular iron measurement

Intracellular iron levels were measured using the FerroOrange (F374, Dojindo) probe according to the manufacturer’s instructions. Briefly, cells were irradiated and incubated for the specified times before being washed with PBS and incubated with a 1 µM FerroOrange solution in the serum-free phenol red-free media (31053028, Gibco) for 30 minutes at 37 °C. Fluorescence (Ex 531/40 – Em 629/53) was directly imaged after the incubation using the Celigo Image Cytometer (Nexcelom).

### Reagents

Ferrostatin-1 (Ferr-1, ab146169-5mg, Abcam) was dissolved in DMSO at 10 mM concentration and diluted in cell culture media to the specified concentrations. Ammonium iron (II) sulfate (09719, Merck) was dissolved in deionized water at 100 mM concentration and diluted in cell culture media to specified concentrations. Deferoxamine (D9533, Sigma Aldrich) was dissolved in DMSO at 100 mM concentration and diluted in cell culture media to the specified concentrations.

### RNA sequencing

RNA samples were extracted from MDA-MB-231 cells 24 hours after being irradiated with 10 Gy in a ^137^Cs irradiator for sequencing as previously described(25). Briefly, RNA was extracted using the RNeasy Mini Kit (74104, Qiagen) following the manufacturer’s protocol. Library preparation, sequencing (paired-end 150 bp sequencing, 20 million reads), and bioinformatic analysis were performed by Novogene Co. (Cambridge, UK) using the Illumina NovaSeq platform. The genes that significantly altered by 10 Gy RT were analyzed by Enrichr (26–28). The Gene Set Enrichment Analysis (GSEA) was performed using gene sets obtained from the Molecular Signatures Database (MSigDB v2024.1.Hs) (29). Raw gene count data were normalized to transcripts per million (TPM) prior to GSEA analysis. Analyses were conducted using the GSEA software (v4.3.3) from the Broad Institute. Gene sets with nominal *P*-value < 0.05 and False Discovery Rate (FDR) < 25% were considered significantly enriched. Additionally, the heatmap was generated using Python (3.11.8).

### Radiation

RT was performed in the Radiation Biophysics facility at the University of Oxford. Cells were exposed to γ-rays from a ^137^Cs sealed source irradiator (GSR D1, Gamma-Service Medical, Leipzig, Germany; dose rate of 1.7 Gy/min). For alpha particle treatment, cells were irradiated with 3.3 MeV alpha particles (LET = 121 keV μm^-1^) using the Oxford ^238^Pu irradiator (30).

### FLASH radiation

Conventional and FLASH RT was performed using an electron linear accelerator which delivers electrons of 6 MeV nominal energy in a circular horizontal beam collimated to 5 cm in diameter, previously described with more details (31, 32). *In vitro* experiments were performed in T12.5 flasks or 35 mm cell culture dishes irradiated one by one in a vertical position in the central part of the beam (Fig. S7). For *in vivo* experiments, the mice were anesthetized with isoflurane supplemented with ∼55% oxygen (90% oxygen evenly mixed with air), then placed in an immobilization cradle and irradiated in an upright position. A 6 mm brass collimator with a 20 mm × 20 mm or a 33 mm × 30 mm aperture was used to further define the RT field covering the whole thorax or the whole abdominal area respectively (Fig. S7B). RT was delivered with 3.5 µs electron pulses with a repetition rate of 25 or 300 Hz (25 Hz for conventional and 300 Hz for FLASH). Dose rates were 0.1 Gy/s for conventional and ≥ 2 kGy/s for FLASH, with a dose-per-pulse of ∼4 mGy and 5 Gy, respectively. Doses ranging from 5 to 20 Gy were given for the *in vivo* and *in vitro* experiments.

The prescribed doses were verified during delivery with pieces of Gafchromic EBT-XD film (Ashland Inc, Covington, KY) placed at the surface or between (abdominal RT) two pieces of Perspex representing a mouse phantom positioned exactly as the mice in the beam path (32). The films were scanned (Epson Perfection v850 Pro, Seiko Epson Corporation, Nagano, Japan), 24 hours post-RT, and the red channel was analyzed for a 20 mm × 20 mm central part of the exposed film, using ImageJ (version 1.52a, Wayne Rasband). The film had previously been calibrated in a clinical 6 MeV electron beam from a Varian Truebeam (Varian Medical Systems Inc, Palo Alto, CA) linear accelerator at the Churchill Hospital site in Oxford, UK. Online verification of the delivered dose was achieved using an Advanced Markus ionization chamber (PTW-Freiburg GmbH, Freiburg, Germany) positioned in the beam (corrected for ion recombination) on the beam energy monitor and collimator system (not disturbing the collimated part of the beam), as well as an upstream positioned toroidal beam charge monitor (33, 34). The beam energy monitor was used to verify that the electron beam energy was consistently 6 MeV, for both FLASH and conventional RT (31). The overall uncertainty in dosimetry was estimated to be 4% (including a measured output variation of within 2%).

### Proton RT

Proton beam RT was given as previously described(35). After placing cell dishes at the midpoint of a 10 cm wide spread-out-Bragg-peak (SOBP, a 230 MeV proton beam generated by a proton therapy system (Sumitomo Heavy Industries, Ltd., Niihama, Japan) was given at a dose rate of 2.14 Gy/min at the Samsung Proton Therapy Center in Seoul, South Korea.

### Lipid peroxidation

Lipid peroxidation was assessed using the BODIPY 581/591 C11 (D3861, Invitrogen) probe according to the manufacturer’s instructions. Briefly, cells were irradiated and incubated for the specified times before incubation with a 2 µM BODIPY 581/591 C11 solution in PBS for 30 minutes at 37 °C. The cells were detached and washed with PBS before FITC fluorescence intensity (oxidized BODIPY emission) was measured using a CytoFLEX (Beckman Coulter) flow cytometer. FlowJo software was used to assess the geometric mean of the intensity of oxidized BODIPY 581/591 C11.

### Clonogenic assay

For ¹³Cs irradiated clonogenic assays, single cells were plated in 6-well plates and allowed to settle for 24 hours before RT. For FLASH vs. conventional irradiated clonogenic assays, cells were seeded in T12.5 flasks and treated with the specified concentrations of Ferr-1 for 24 hours before RT. Prior to RT, the flasks were positioned vertically. Following RT, the cells were detached, counted, and replated as single cells in 6-well plates. Seeding densities were optimized for each cell line and RT dose. Colonies were cultured for 8 to 12 days before being stained with a 20% methanol/0.5% crystal violet solution. Colonies were then counted to determine plating efficiency (PE = colony number/number of cells plated), and the surviving fraction (SF) was calculated using the formula: SF = (number of colonies formed after treatment) / (number of cells seeded × PE), following the methodology of a previous study (36).

### Patient data

In total, 748 genes in the Cancer Gene Consensus were acquired from the Catalogue Of Somatic Mutations In Cancer (COSMIC) and analyzed by DAVID functional analysis to identify the enriched pathways (37, 38). Tissue microarrays were purchased from Biomax (BR1008b – Breast cancer, LC10011b – Lung cancer). *TFRC* expression data from the TCGA PanCancer Atlas was obtained from cBioPortal, including mRNA expression z-scores (log RNA Seq V2 RSEM) normalized relative to normal samples, along with corresponding clinical data (39). We have stratified “high TFRC” group as patients with higher TFRC expression z-score compared to normal tissue samples and “low TFRC” group as patients with lower TFRC expression z-score compared to normal tissue samples. *TFRC* expression across different tumor types was visualized using TCGA datasets through The University of ALabama at Birmingham CANcer data analysis Portal (UALCAN) (40).

### In vivo experiments

All animal experiments were performed according to the guidelines of the United Kingdom Home Office and the University of Oxford under the project licenses PP8415318 and PP4558762. Female BALB/cAnNCrl mice were purchased from Charles River (6-8 weeks, strain code 028) and housed in individually ventilated cages containing no more than six mice per cage, in a 12/12-hour light/dark cycle. For normal mouse lung studies, lung tissues were collected 24 hours or 7 days after RT either at conventional or FLASH dose rates. To determine iron accumulation in the upper intestine, mice were fed with either control diet with 200 ppm iron (TD. 08713, Envigo) or a high-iron diet with 5000 ppm iron (TD140464, Envigo) for 24, and 48 hours (41). Intestinal tissues were collected at the specified time points to assess the correlation between iron diet and RT effects. For RT studies, mice were fed either a control diet or a high-iron diet for evaluation. After 24 hours, they were either kept as controls or exposed to conventional or FLASH dose rate RT. Post-RT, mice were monitored and weighed daily. One group of mice returned to a control diet, and the other group of mice remained on a high-iron diet. Seventy-two hours after RT, mice were sacrificed to collect blood samples and intestinal tissues.

For Ferr-1 treatment (341494-25mg, Merck), mice received intraperitoneal injection of 2 mg/kg of Ferr-1 one day before RT and then once daily until mice were sacrificed 72 hours after RT.

For tissue processing, lung tissues were inflated with 10% formalin and kept in formalin for 24 hours before storing in 70% ethanol. Intestinal samples were flushed with PBS, fixed in formalin for 24 hours. Then they were cut with micro scissors to expose the lumen. The tissues were then rolled from the posterior end to reveal the inner lumen and stored in 70% ethanol until being processed and embedded in paraffin. Sections of 6 µm thickness were prepared for immunohistochemistry analysis using microtome.

### Mouse tumor tissues

A549 and Calu-6 xenograft tumor tissues were treated with conventional, or FLASH RT. Briefly, A549 and Calu-6 cells were subcutaneously injected into athymic nude mice. When the tumor size reached 100 mm^3^, mice were randomized into three groups: control, conventional RT, and FLASH RT (15 Gy for A549, 20 Gy for Calu6). When the tumor size reached 800 mm³ or 50 days post-RT, the mice were sacrificed to collect tumor tissues. The harvested tissues were fixed in formalin, then stored in 70% ethanol until further processing and paraffin embedding. Paraffin-embedded tissues were cut into 6 µm sections.

### Immunohistochemistry

Paraffin-embedded tissues were sectioned and stained as previously described (25, 42). Briefly, after incubation in a 65 °C oven for 30 minutes, slides were deparaffinized by immersing in Histo-Clear (National Diagnostics, HS-200) twice for 3 minutes followed by rehydration for 3 minutes in 100 %, 70 %, and 50 % ethanol. The slides were washed in deionized water for 5 minutes before antigen retrieval was performed in 1 x Citrate buffer (10 x Citrate buffer (pH 6.0), C9999, Sigma; diluted with deionized water and 0.05 % Tween-20) which was heated to 110 °C in a pressure cooker for 3 minutes. Then, the slides were cooled at room temperature for 20 minutes. Endogenous peroxidase and phosphatase activities were blocked by Dual Endogenous Enzyme Block (S2003, Agilent Technologies) for 20 minutes. The slides were blocked with 10 % goat serum for 30 minutes and incubated with antibodies (anti-4-HNE antibody (BS-6313R, Thermo Fisher or ab48506, abcam) diluted in 1:100 or 1:200 with antibody diluent (ab64211, abcam) or anti-OLFM4 antibody (39141, Cell Signaling) diluted in 1:300 with antibody diluent) at 4 °C overnight. After washing three times with PBS/0.05 % Tween-20, the secondary antibody with HRP-labelled polymers was added to slides (K4003 or K4001, Agilent Technologies) and incubated at room temperature for 20 minutes. Slides were then washed three times with PBS/0.05% Tween-20 and incubated with DAB (3,3′-Diaminobenzidine) (K3468, Agilent Technologies) to visualize the HRP signals. The slides were counterstained with hematoxylin (1.09249.0500, Sigma Aldrich) and mounted with aqueous mounting medium (1.08562.0050, Merck).

Tissue iron was stained using a Prussian Blue staining kit (ab150674, Abcam) according to the manufacturer’s protocol, with a peroxide-DAB enhancing step. Briefly, slides were deparaffinized and incubated with Prussian Blue solution for 1 hour at room temperature before washing 3 times with deionized water. If no blue prussiate was visible, slides were incubated with a 0.3% hydrogen peroxide and 0.01M sodium azide methanol solution for 30 minutes at room temperature. Slides were then washed 4 times with PBS, incubated with DAB until tissues turned brown. Slides were washed with 3 changes of deionized water before counterstaining with nuclear fast red for 4 minutes at room temperature. Slides were dehydrated in 50%, 70%, and 100% ethanol and 2 changes of 100% xylene before mounting with DPX (06522, Sigma Aldrich). TUNEL staining was performed following manufacturer’s protocol (C10625, Thermo Fisher).

Hematoxylin and eosin (H&E) staining was performed by deparaffinizing tissues followed by incubation with 25% Harris’ hematoxylin (CODE) diluted in water for 3 minutes. Slides were dipped 10 times in acid alcohol (1% hydrochloric acid in 70% ethanol) and then washed with 10 dips in tap water. Slides were dipped 10 times into Scott’s tap water and then washed with 10 dips in tap water. Slides were dipped 10 times in 0.5% lithium carbonate and then washed with running tap water. Slides were then dipped 10 times in 50%, 70%, and 90% ethanol. Slides were then incubated in alcoholic eosin for 1 minute before being dipped 10 times 90% ethanol and repeated with fresh 90% ethanol. Slides were then dehydrated in 100% ethanol and 2 changes of 100% xylene before mounting with DPX.

All slides were imaged with an Aperio Scanscope CS digital pathology scanner (Leica Biosystems Imaging, Inc.) at 40X or 20X magnification.

For 4-HNE staining, three representative fields were imaged per slide at 10x or 20 x magnification. DAB staining intensity was analyzed using FIJI (Image-J-based open-source software) (43). Images were chosen from the irradiated lung lobe and upper intestine areas where iron is mainly accumulated. Positively stained areas were extracted using color deconvolution with the brightness and contrast optimized for each set of images and the values were kept consistent across all images from the same set. Positively stained areas were converted into binary images, and the integrated density values were calculated.

The amount of iron in the upper intestines or microarray cores were assessed using FIJI. First, positively stained iron areas were selected and extracted by using color threshold, converted into binary images and positive areas were measured. Similarly, total tissue areas stained with nuclear fast red were selected and extracted using the color threshold, converted into binary and measured. The percentage of iron in each tissue relative to total cellularity was determined using the iron positive and total tissue areas.

For TUNEL staining, three representative fields at 10x magnification were selected per slide. Positive cell detection was conducted using QuPath software (version 0.5.0) built-in algorithm, with optimized parameters for cell detection and signal intensity to reliably identify TUNEL-positive nuclei. Detection thresholds, cell size estimates, and background subtraction settings were adjusted to maximize accuracy and consistency across samples. The percentage of TUNEL-positive cells was calculated by dividing the number of TUNEL-positive nuclei by the total number of detected nuclei within each region of interest.

Three representative fields of H&E stained images 1 mm in size were randomized and the remaining crypts were manually counted (Fig. S8). The percentage of remaining crypts was determined by comparing to the untreated and non-irradiated control. The number of OLFM4 positive crypts were manually counted in three representative fields per slide, each within a 1 mm area.

### Ferritin ELISA

Mouse serum was collected by centrifugation of mouse blood at 3000 rpm for 10 minutes. Serum ferritin levels were measured by using a commercially available mouse ferritin ELISA kit according to the manufacturer’s protocol (Abcam, #ab157713).

### Statistical analysis

The data are presented as the mean, the standard deviation, and the individual observations of at least three replicates. The data were analyzed using GraphPad Prism software (GraphPad Prism 10, Dotmatics) as indicated in each figure. Briefly, for comparisons between two groups, we performed two-tailed Student’s *t*-tests. Nonparametric Kolmogorov-Smirnov test was used for tissue microarray data. To compare multiple groups, we used one-way ANOVA for single-condition experiments and two-way ANOVA for experiments involving multiple conditions. A *P* value of <0.05 was considered statistically significant.

### Study approval

This study involved mouse studies. Experimental procedures were carried out under project licenses (PP8415318 and PP4558762) issued by the UK Home Office under the UK Animals (Scientific Procedures) Act of 1986.

## Results

### RT induces lipid peroxidation and ferroptosis

RT induces cell death primarily through the induction of DNA damage. Recently, ferroptosis has been identified as a contributor to RT-induced cell death that occurs independently of DNA damage (7–10). RNA sequencing was performed in MDA-MB-231, a human breast cancer cell line, treated with 0 or 10 Gy of RT to determine changes in gene expression induced by RT. Cells were irradiated with γ-rays from a ^137^Cs irradiator 24 hours before RNA extraction and transcriptomic analysis. Kyoto Encyclopedia of Genes and Genomes (KEGG) enrichment analysis using Enrichr identified significant changes in genes associated with the cell cycle, DNA replication, and p53 pathway, which are hallmarks of cells exposed to RT-induced DNA damage (26–28) (Fig. 1A). Additionally, RT also affected genes involved in ferroptosis (Fig. 1A, Fig. S1A) in line with previous studies (7–9). Gene Set Enrichment Analysis (GSEA) further identified a strong association between RT exposure and significant alterations in ferroptosis-related genes (29) (Fig. 1B, Fig. S1B). To assess the biological significance of the changes in gene expression, the effect of RT on lipid peroxidation and ferroptosis was examined in MDA-MB-231 (human breast cancer) and A549 (human lung cancer) cell lines *in vitro*. We first detected lipid peroxidation 24 hours after exposure to 4 Gy of RT, with levels increasing in a dose-dependent manner up to 10 Gy (Fig. 1C and 1D). There was also a time-dependent increase in lipid peroxidation when irradiating cells at 10 Gy with a statistically significant enhancement detected as early as 4 hours post RT (Fig. 1E and 1F) and a further progressive increase up to 72 hours (Fig. 1G and 1H). In addition, we also investigated how different RT sources affected RT-induced lipid peroxidation using alpha particles and protons, indicating that various RT modalities increased lipid peroxidation (Fig. S2). Since increases in lipid peroxidation can lead to ferroptosis, clonogenic survival was assessed using ferrostatin-1 (Ferr-1), a selective inhibitor of ferroptosis. We found that Ferr-1 significantly enhanced clonogenic survival in both cell lines following RT, confirming that RT-induced ferroptosis contributes to the decreased survival of these tumor cells (Fig. 1I and 1J).

**Fig 1.**
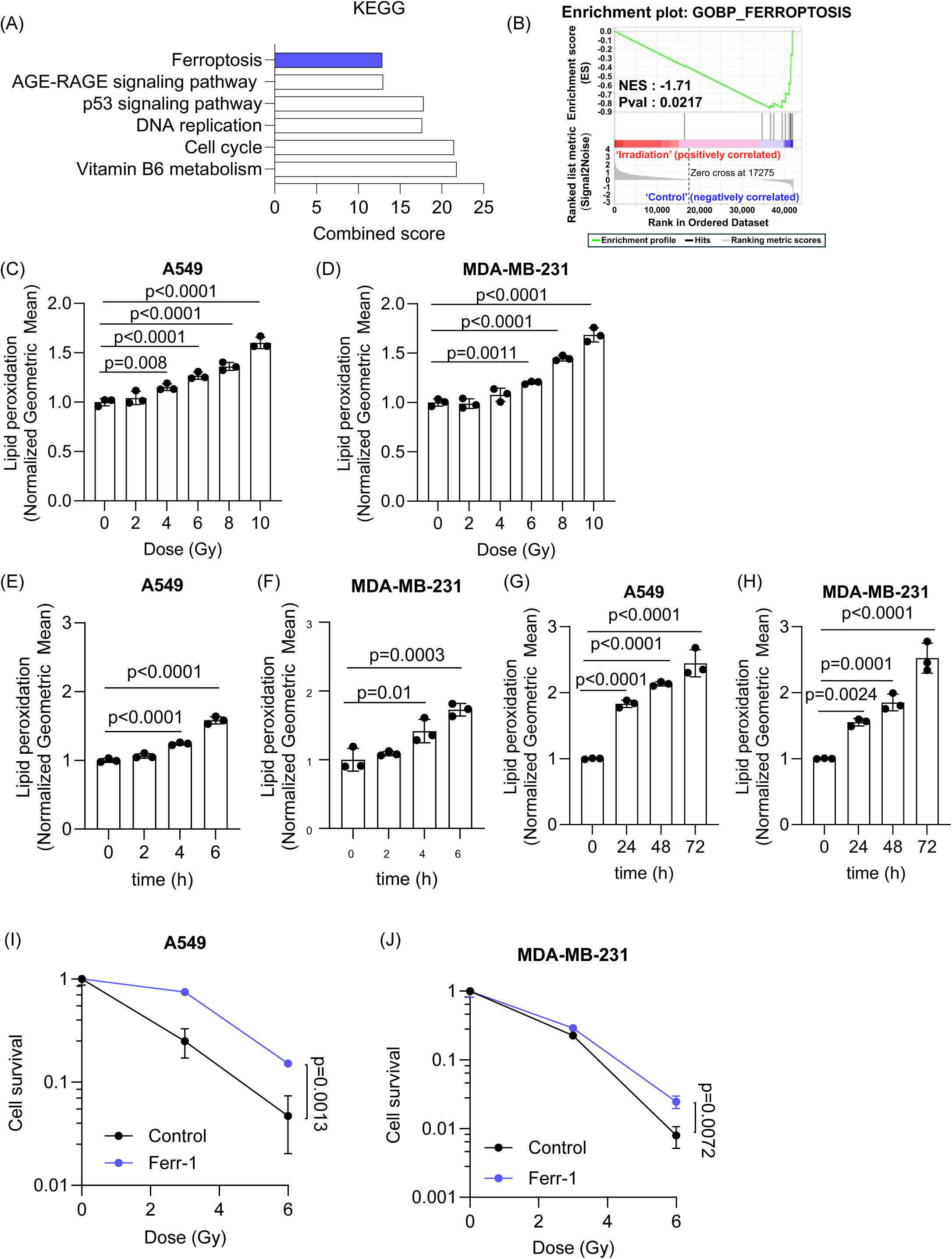
RT induces lipid peroxidation and ferroptosis in cancer cells in a dose and time-dependent manner. (A) RNA sequencing was performed after treating MDA-MB-231 cells with 0 or 10 Gy of RT. Pathways analysis by Enrichr identified that genes involved in the “Cell Cycle”, “p53 signaling pathway”, “DNA replication” and “Ferroptosis” were significantly altered by RT. (B) GSEA enrichment further confirmed significant alterations of genes involved in ferroptosis after RT. (C-H) Lipid peroxidation was measured using C11 BODIPY in A549 and MDA-MB-231 24 hours after RT at varying doses from 2 to 10 Gy (C-D) as well as at different time points after 10 Gy of RT (E-H). (I-J) Clonogenic survival was determined after RT with or without pre-treatment of 20 µM (A549, I) or 4 µM (MDA-MB-231, J) Ferr-1. RT was given using a ^137^Cs irradiator. Error bars indicate standard deviation (SD) (n = 3 per group). Statistical tests were performed by One-way ANOVA with Dunnett’s multiple comparisons test (C-H) and unpaired *t*-test (I-J).

### FLASH RT induces ferroptosis in tumor cells

FLASH, an ultra-high dose rate RT, has been shown to result in less lipid peroxidation in micelles and liposomes (22). To investigate whether FLASH RT also reduces lipid peroxidation in tumor cells, A549 and MDA-MB-231 were treated with electrons at conventional (6 MeV, 0.1 Gy/s) or FLASH RT (6 MeV, > 2 kGy/s) dose rates. The RT dose used in these experiments was 10 Gy, as the FLASH effect is larger at higher RT doses (≥ 10 Gy) (15, 16, 44, 45). Interestingly, FLASH RT induced lipid peroxidation in both cancer cell lines, and it was comparable to levels found with conventional RT (Fig. 2A and 2B). Furthermore, tumor cells treated with doses of RT ranging from 5 to 20 Gy exhibited a dose-dependent increase in lipid peroxidation both by FLASH RT and conventional RT dose rates (Fig, 2C and 2D). Clonogenic assays using Ferr-1 confirmed that Ferr-1 increased cell clonogenicity in both conventional and FLASH RT treated cells, indicating that ferroptosis contributes to cell death after conventional and FLASH RT in cancer cells (Fig. 2E and 2F).

**Fig 2.**
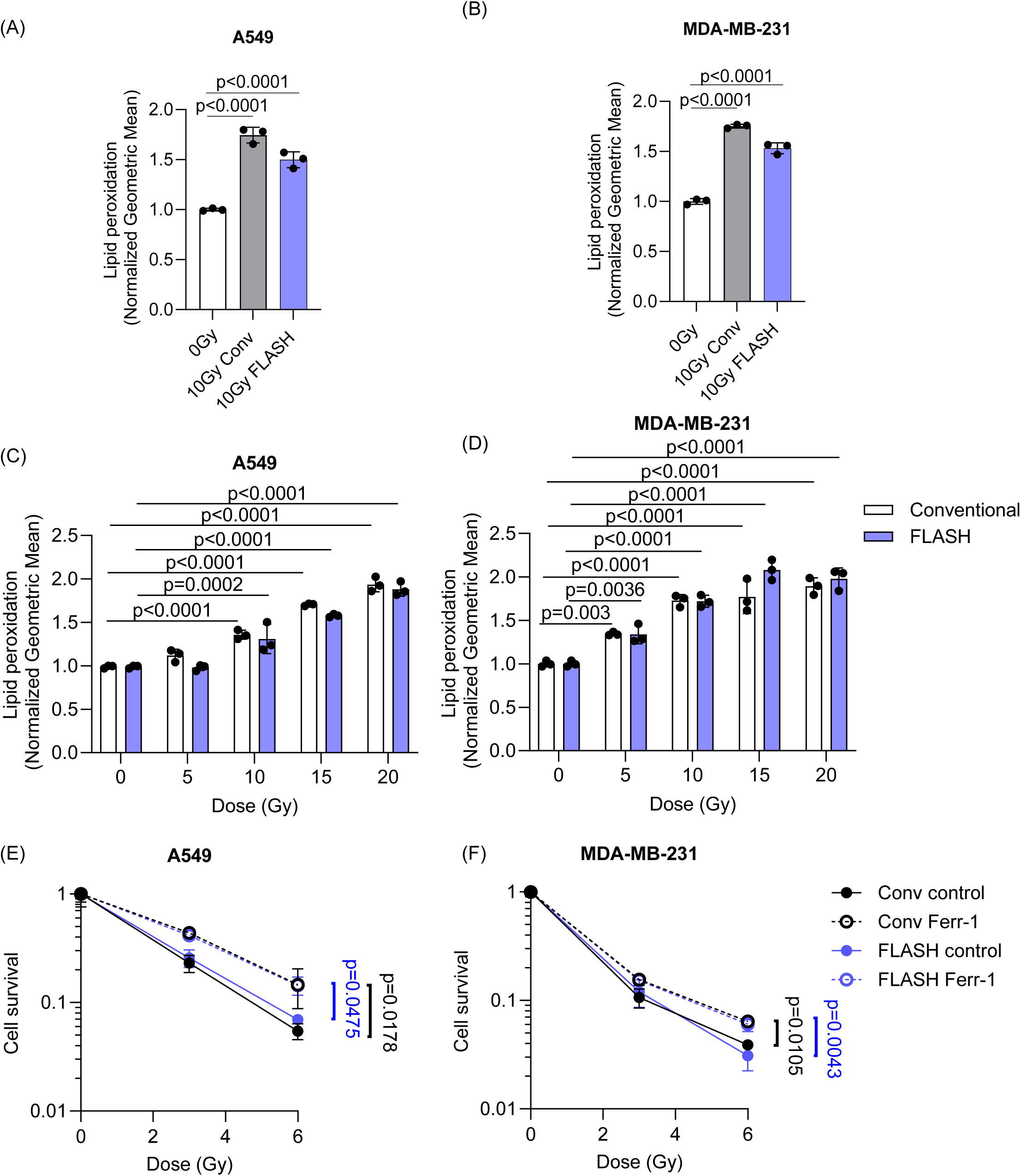
FLASH RT induces similar levels of lipid peroxidation and ferroptosis with conventional RT in cancer cells. (A-B) A549 (A) and MDA-MB-231 (B) cells were irradiated at 10 Gy with conventional or FLASH dose rates with an electron linear accelerator. A significant increase in lipid peroxidation was observed 24 hours after RT using C11 BODIPY in all treatment groups. (C-D) A significant dose-dependent induction of lipid peroxidation was observed after conventional and FLASH RT. (E-F) Clonogenic survival was determined after conventional or FLASH RT with or without pre-treatment of 20 µM (A549) or 4 µM (MDA-MB-231) Ferr-1. Error bars indicate standard deviation (SD) (n = 3 per group). Statistical tests were performed by One-way ANOVA with Dunnett’s multiple comparisons test (A-B), Two-way ANOVA with Tukey’s multiple comparisons test (C-D), and One-way ANOVA with Tukey’s multiple comparisons test (E-F).

### FLASH RT induces lipid peroxidation in tumors but not in normal tissues

The tumor microenvironment consists of complex interactions between the tumor and its surrounding counterparts, including fibroblasts, blood vessels, and low oxygen levels (46). To determine whether the effect of FLASH RT on lipid peroxidation could be detected *in vivo*, tumor tissues were collected from mouse xenograft models of two different human lung cancer cell lines (A549 and Calu6) for staining with 4-hydroxynonenal (4-HNE), a classical marker for lipid peroxidation. Consistent with the *in vitro* observations, both conventional and FLASH RT (15 Gy or 20 Gy) increased 4-HNE staining in these tumor tissues compared to the non-treated control group (Fig. 3A).

**Fig 3.**
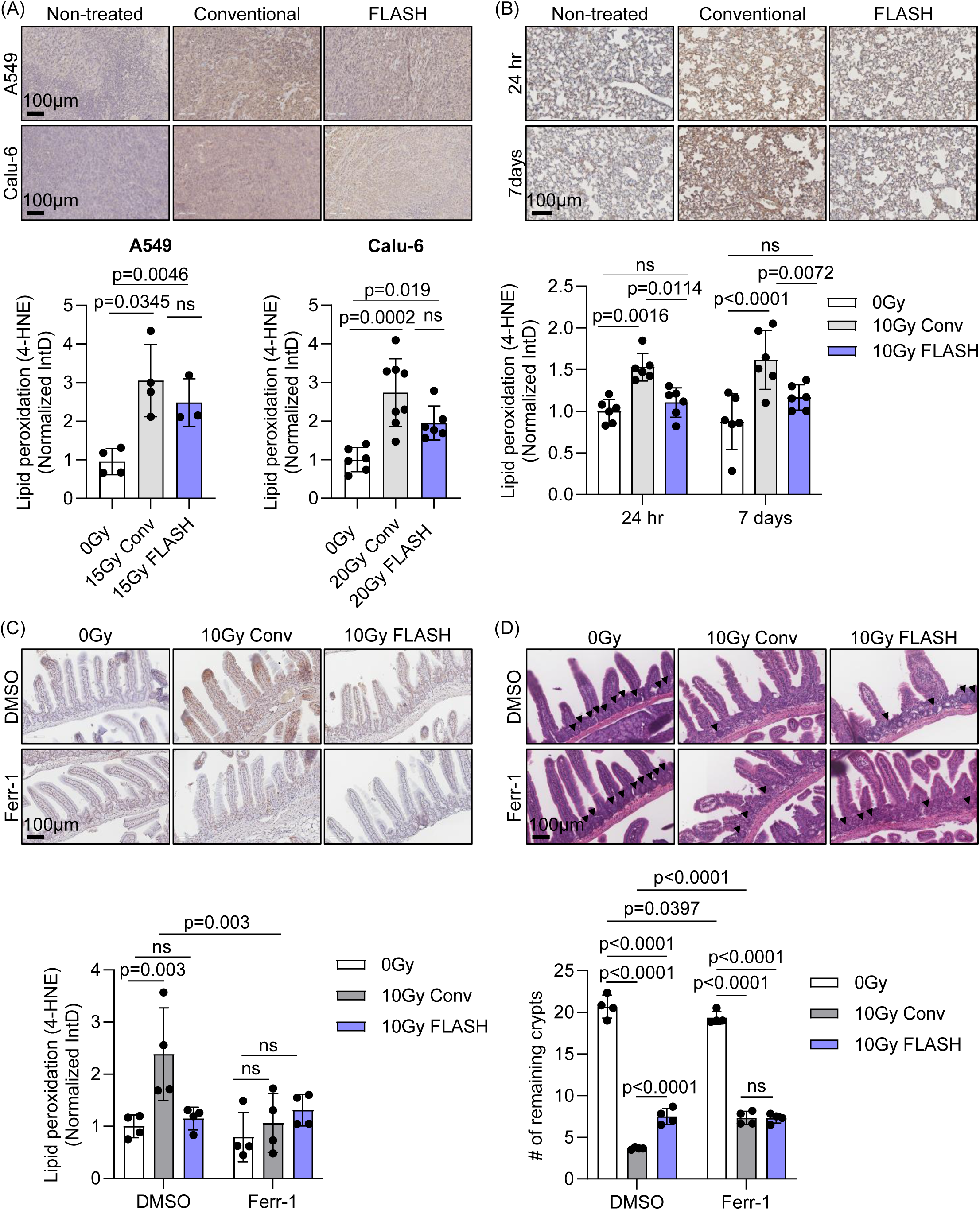
FLASH RT induces lipid peroxidation in mouse xenograft tumors while sparing normal lung and intestine tissues. (A) Lung tumor tissues (A549 and Calu6) from subcutaneous mouse xenograft models were stained with 4-HNE, a lipid peroxidation marker. After conventional and FLASH RT (15 Gy for A549 and 20 Gy for Calu6), there was a significant increase in 4-HNE staining intensity (n = 4 for A549 0 Gy and conventional RT, n = 3 for FLASH RT, n = 6 for Calu-6 0 Gy and FLASH RT, n = 8 for Calu-6 conventional RT). (B) BALB/c mouse lung tissues were stained with 4-HNE after 10 Gy of conventional or FLASH RT. Lipid peroxidation was significantly enhanced 24 hours and 7 days after conventional RT. However, FLASH RT did not change lipid peroxidation (n = 6 per group at each time point). (C-D) BALB/c mice were treated daily with 2 mg/kg Ferr-1 or DMSO vehicle, starting one day prior to RT, to inhibit ferroptosis. Lipid peroxidation, detected by 4-HNE staining, was markedly increased after 10 Gy conventional RT compared to 0 Gy, but not after 10 Gy FLASH RT in the DMSO-treated group (C). This increase in lipid peroxidation was reversed by Ferr-1 treatment. Tissue damage in the upper intestines was assessed by H&E staining, based on the number of remaining intestinal crypts (D, arrowheads). Both 10 Gy conventional and FLASH RT significantly reduced crypt numbers, with conventional RT causing more severe damage. Ferr-1 treatment improved crypt preservation in the conventional RT group but had no significant effect in the 0 Gy or FLASH RT groups (n = 4/group). Error bars indicate standard deviation (SD). Statistical tests were performed by One-way ANOVA (A) or Two-way ANOVA (B-D) with Tukey’s multiple comparisons test. IntD: Integrated Density.

In contrast to tumors, we did not observe similar effects between conventional and FLASH RT in normal mouse lung tissues, supporting previous observation of reduced lipid peroxidation (22, 24). Conventional RT treatment of the whole thorax of BALB/c mice at 10 Gy significantly increased lipid peroxidation in lung tissues at 24 hours and 7 days post-treatment. In contrast, the same dose of FLASH RT resulted in markedly decreased lipid peroxidation compared to conventional RT across all tested conditions. (Fig. 3B). Additionally, using TUNEL assay, a reduction in apoptosis was observed in lung tissues after FLASH RT compared to conventional RT, indicating decreased tissue damage (Fig. S3).

In mouse intestines, FLASH RT at 10 Gy also led to reduced lipid peroxidation as well as crypt damage determined by the number of remaining crypts relative to conventional RT (Fig. 3C and 3D).

Interestingly, treatment with Ferr-1 also decreased lipid peroxidation and crypt damage to levels comparable to those observed with FLASH RT. These findings suggest that reduced lipid peroxidation levels induced by FLASH RT in normal tissue compared to tumors may contribute to the reduced normal tissue damage observed with FLASH RT through ferroptosis.

### Iron is essential for tumor survival

Iron is an essential element to maintaining cellular functions such as oxygen transport, energy production, hematopoiesis as well as DNA replication (1). Cancer cells strongly depend on iron for survival and as a result, increased levels of iron and ferritin, an iron storage protein, are often observed in tissues and serum of various cancer patients (2–5). Iron staining in tissue microarrays (TMA) of breast and lung cancer patients using Prussian Blue with DAB (3,3′-Diaminobenzidine) indicated that iron was highly elevated in both lung (Fig. 4A) and breast (Fig. 4B) cancer samples compared to corresponding normal tissues. The functional enrichment analysis of 748 genes from the Catalogue of Somatic Mutations in Cancer (COSMIC) Cancer Gene Consensus further confirmed that cancer driving genes involved in iron metabolism or homeostasis including “4 iron 4 sulfur cluster binding”, “response to iron ion”, and “cellular iron ion binding” were significantly amplified, supporting the crucial needs of iron in cancer (Fig. 4C) (37). Specifically, transferrin receptor (TFRC), a receptor facilitating iron transport, was included in COSMIC. Expression of TFRC was significantly correlated with poor overall survival in all cancer types (Fig. 4D) and its expression was highly upregulated in the majority of cancer types (Fig. 4E). The inhibition of *TFRC*, which inhibited iron uptake, significantly decreased cell survival in both lung (A549) (Fig. 4F and Fig. S4) and breast (MDA-MB-231) cancer cells (Fig. 4G), indicating that iron is critical for tumor cell survival.

**Fig 4.**
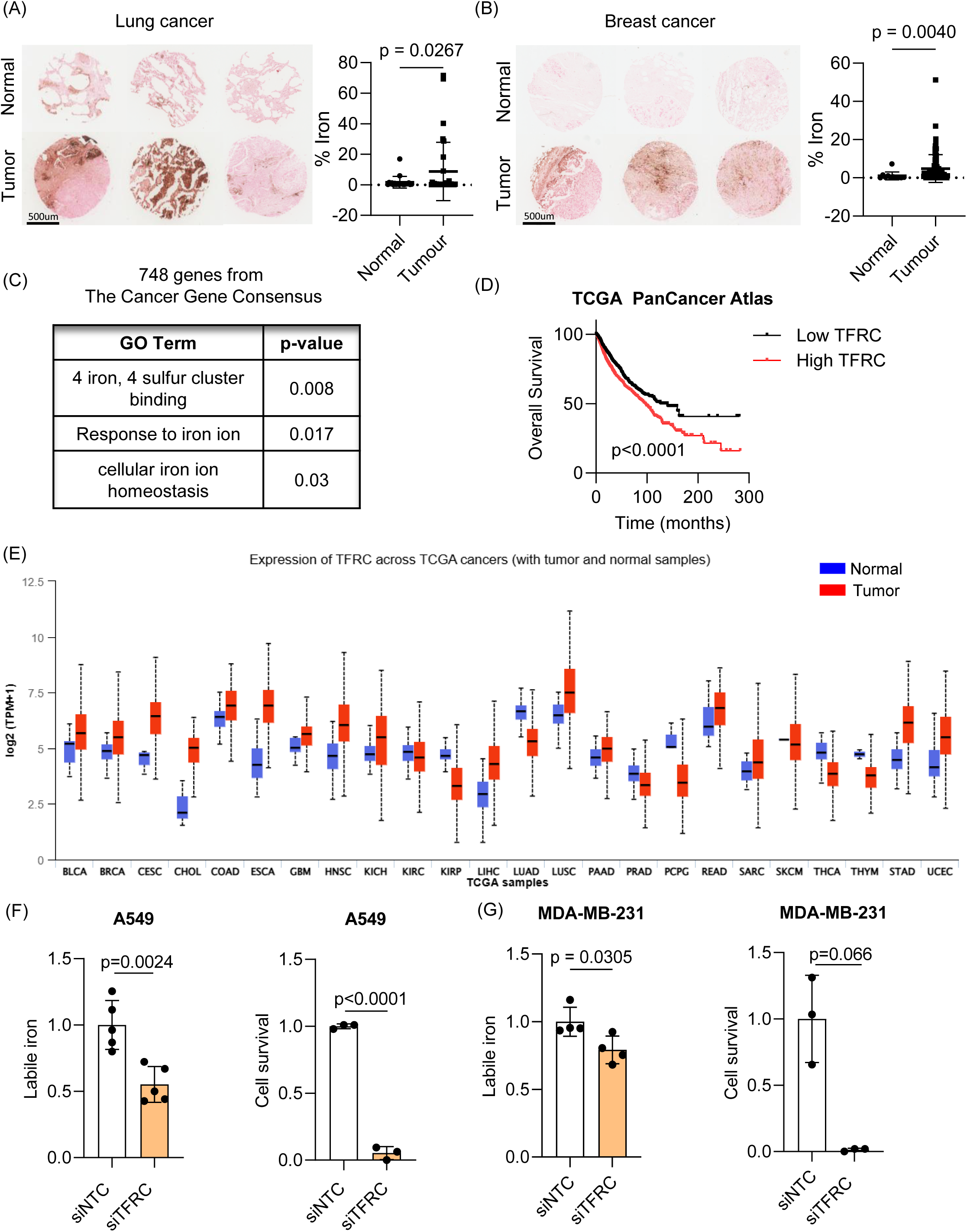
Iron is essential for tumor survival. (A-B) Tissue microarray slides from breast and lung cancer patients were analyzed to measure iron levels using Prussian blue. Compared to normal lung (A) or breast tissues (B), iron levels were significantly higher in lung and breast cancer tissues (Lung cancer; n = 20 for normal tissues, n = 32 for lung adenocarcinoma tissues, Breast cancer; n = 11 for normal tissues, n = 90 for cancer tissues). (C) The functional enrichment analysis of 748 genes from the Catalogue of Somatic Mutations in Cancer (COSMIC) Cancer Gene Consensus showed significantly regulated pathways of cancer driving genes “4 iron 4 sulfur cluster binding”, “response to iron ion”, and “cellular iron ion binding”. (D) Analysis of TCGA PanCancer Atlas data revealed that cancer patients exhibiting higher *TFRC* expression in tumor tissues compared to normal tissues had significantly poorer overall survival outcomes (Low *TFRC*: n=2014, High *TFRC*: n=3257; log-rank test, p < 0.0001) (E) TCGA data showing *TFRC* mRNA expression in normal and various cancer patient tissues were visualized using UALCAN (F-G) The role of iron in tumor cell survival was determined by knocking down *TFRC*, an iron transporter. Compared to the siRNA against scrambled sequence, the inhibition of *TFRC* expression significantly decreased intracellular iron and cell survival measured by clonogenic assay in both A549 (F) and MDA-MB-231 (G) cells. Statistical tests were performed by nonparametric Kolmogorov-Smirnov test (A-B), log-rank (C), and upaired *t*-test (F-G).

### Iron enhances RT sensitivity and ferroptosis

Since iron is central to lipid peroxidation, increasing iron levels by adding ammonium iron (II) sulfate also increased lipid peroxidation significantly after 10 Gy RT in both A549 (Fig. 5A) and MDA-MB-231 (Fig. 5B). The treatment of Ferr-1 abrogated the increased lipid peroxidation induced by iron. These results were supported by decreased clonogenic survival upon ammonium iron (II) sulfate treatment. The radiosensitizing effect of iron was rescued by Ferr-1 treatment, suggesting that iron supplementation indeed leads to RT-induced ferroptosis (Fig. 5C and 5D). Consistent with a previous report (9), iron depletion using deferoxamine (DFO), an iron chelator, also reduced lipid peroxidation (Fig. 5E and 5F) and RT-induced tumor cell death (Fig. 5G and 5H). Overall, these data highlight the critical role that iron plays in tumor cell survival and ferroptosis in response to RT.

**Fig 5.**
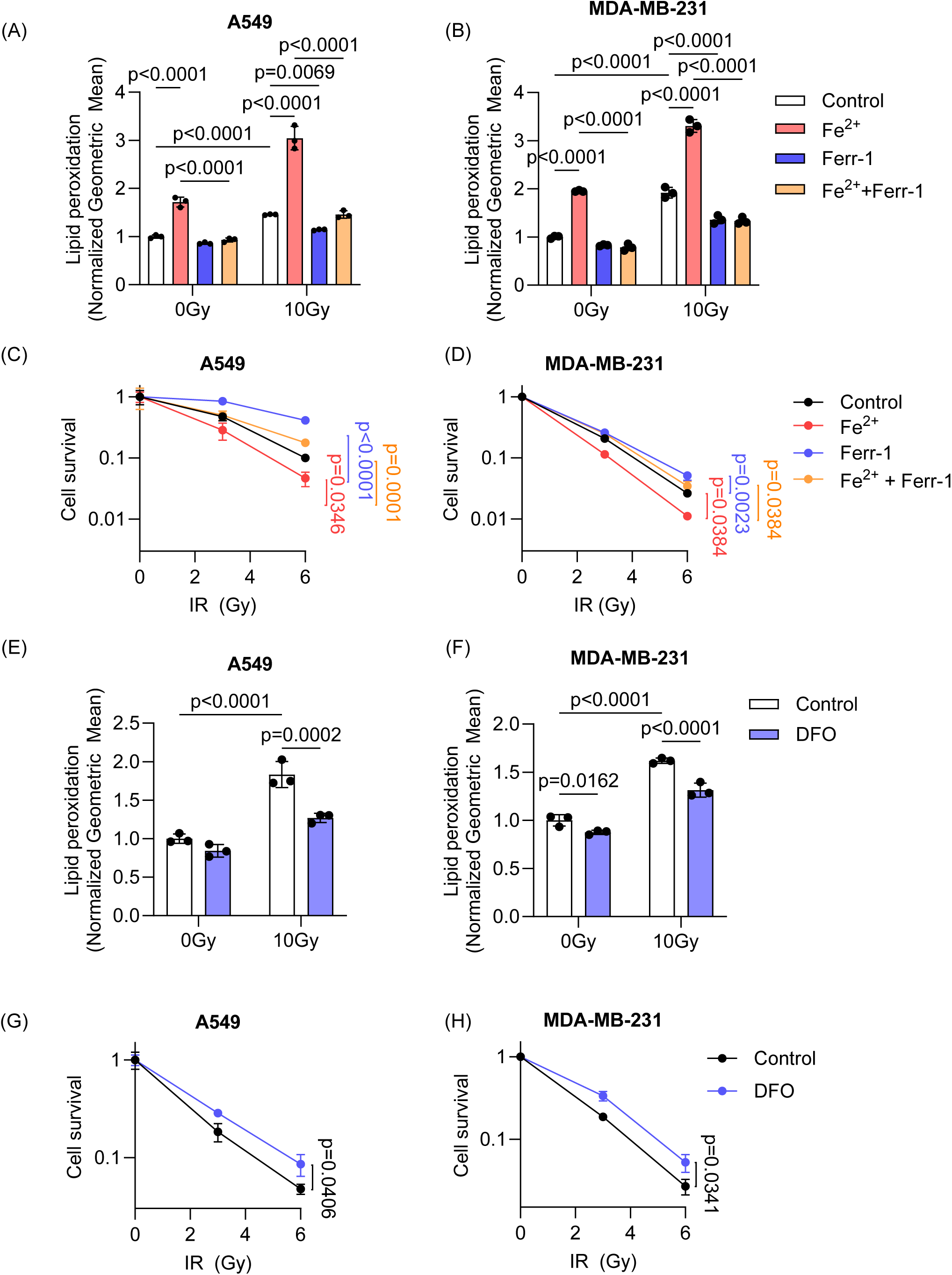
Iron enhances RT sensitivity and ferroptosis. The role of iron in lipid peroxidation and cell survival was determined by measuring lipid peroxidation using C11 BODIPY and clonogenic assay, respectively. Adding a suboptimal dose of iron (50 μM and 5 μM of ammonium iron (II) sulfate in A549 and MDA-MB-231 respectively) increased lipid peroxidation (A-B) and radiosensitization (C-D) in both cell lines. Treatment with Ferr-1 (20 μM and 4 μM in A549 and MDA-MB-231, respectively) decreased lipid peroxidation induced by RT (A-B) and reversed radiosensitization (C-D) indicating that ferroptosis was further induced by external iron. (E-H) Inhibition of iron availability using deferoxamine (DFO) (200 nM and 100 nM in A549 and MDA-MB-231, respectively) decreased lipid peroxidation induced by RT (E-F) and increased radioresistance (G-H) of A549 and MDA-MB-231 cells. (A-H, n = 3/group) Error bars indicate standard deviation (SD). Two-way ANOVA with Tukey’s multiple comparisons test (A-B), One-way ANOVA with Tukey’s multiple comparisons test (C-D, G-H), and unpaired *t*-test (E-F). For C-D and G-H, 6 Gy conditions were compared.

### Increasing iron levels in normal tissue abolishes FLASH sparing effect

To investigate why the reduction in lipid peroxidation observed with FLASH RT is not evident in tumour tissues, we hypothesized that intrinsic differences in iron levels between normal and tumour tissues may account for this variation. in lipid peroxidation found after FLASH RT compared to conventional RT (23). To test our hypothesis, we increased iron levels specifically in the normal intestine of BALB/c mice. The intestinal tissue is highly sensitive to RT-induced toxicities due to the presence of rapidly dividing cells, particularly in the crypts and endothelium. However, multiple studies have shown that FLASH RT significantly reduces intestinal damage compared to conventional RT (32, 47, 48). Since iron absorption primarily occurs in the small intestine, particularly in the duodenum and proximal jejunum, mice were fed a high-iron diet (5000 ppm iron) to elevate iron levels locally without disrupting systemic iron homeostasis, which is tightly regulated by the hepicidin-ferroportin axis (49). Iron accumulation in the small intestine was confirmed as early as 24 and 48 hours after initiating the high-iron diet (Fig. S5A).

To examine whether high iron levels at the time of RT affect RT responses, mice were fed with a high-iron diet (HI) for 24 hours and treated with RT at conventional or FLASH RT dose rates, then returned to the control diet (HI 24-hour) or remained on a high-iron diet for an additional 72 hours (HI 96-hour) to determine the effect of iron before or after the RT (Fig. 6A). Serum ferritin levels suggested that the iron diet did not cause any systemic changes in iron levels (Fig. S5B). Consistent with our previous observations (Fig. 3C), conventional RT at both 8 and 10 Gy increased lipid peroxidation (4-HNE), with less induction with FLASH RT in the control diet group (200 ppm iron) (Fig. 6B and 6C). Similarly, lipid peroxidation was not significantly different after FLASH RT in mice fed an HI for 24 hours compared to non-irradiated mice, while conventional RT still showed high induction. However, the HI diet before and after RT (HI 96-hour) abrogated the FLASH RT effect with a significant increase in lipid peroxidation relative to the non-irradiated group and showed no significant difference from the conventional RT group.

**Fig 6.**
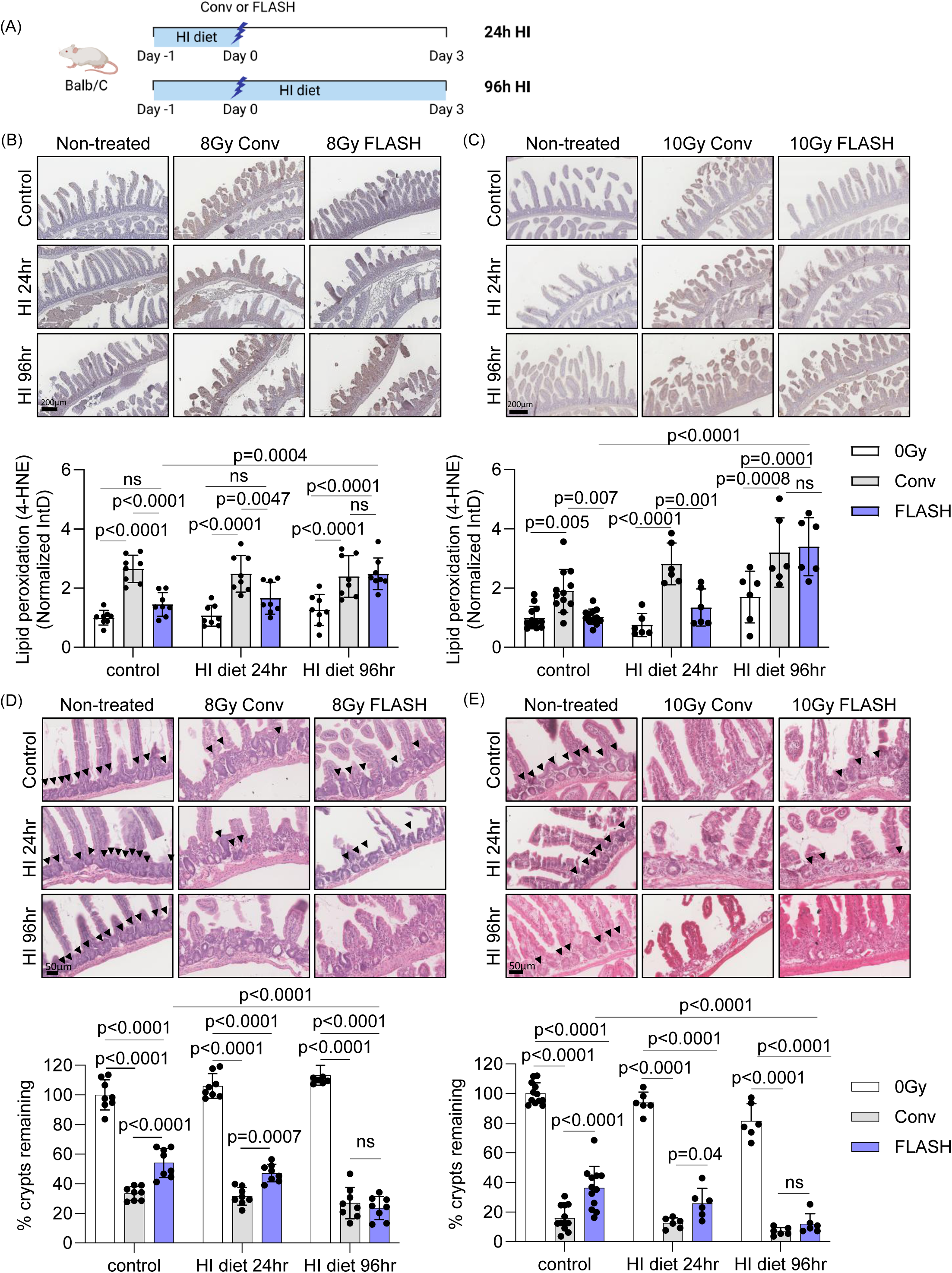
Increasing iron levels in normal tissue reverts FLASH sparing effect. (A) BALB/c mice were fed with high iron diet (5000 ppm iron) for 24 hours to increase iron levels in normal intestine. After RT treatment at 8 or 10 Gy, mice were returned to a normal diet or stayed on a high-iron diet for another 72 hours. (HI: High iron diet) (B-C) Lipid peroxidation, which was stained with 4-HNE was highly increased by both 8 (B) and 10 Gy (C) conventional RT in the control or high iron diet groups both for 24 hours and 96 hours. FLASH RT at 8 or 10 Gy did not increase lipid peroxidation as significantly as conventional RT with the control or 24-hour high-iron diet. However, a prolonged high-iron diet even after RT treatment eliminated the FLASH effect on lipid peroxidation. (D-E) Tissue damage in the upper intestines was determined by H&E staining. The presence of remaining intestinal crypts (arrow heads) was counted and analyzed. The number of remaining crypts was significantly decreased after 8 (D) and 10 Gy of conventional RT in the control or high-iron diet groups for 24 hours and 96 hours. FLASH RT-induced damages were significantly lower than conventional RT in the control or 24-hour high-iron diet mice. However, a prolonged high iron diet, diminished the sparing observed in FLASH irradiated tissues since FLASH RT induced similar tissue damage to conventional RT. (B - E, for 8 Gy, n = 8 in every condition; for 10 Gy, control diet n = 12, HI diet 24 hours or 96 hours n = 6) Error bars indicate standard deviation (SD). Statistical test was performed by Two-way ANOVA with Tukey’s multiple comparisons. IntD: Integrated Density.

These results were further supported by quantifying the number of crypts after RT and FLASH RT (Fig. 6D and 6E). In control diet group, the conventional RT groups exhibited a significantly lower number of remaining crypts compared to the FLASH RT groups following treatment either at 8 or 10 Gy. A high-iron diet for 24 hours (HI 24-hour) did not alter this trend of intestinal damage. Notably, in the HI 96-hour mice, the difference between conventional and FLASH RT groups was no longer significant. The prolonged high iron diet exposure resulted in a reduced number of surviving crypts even in the FLASH RT groups, eliminating the FLASH sparing effect observed in other conditions. These findings were further supported by OLFM4 staining, which was used to assess the number of remaining intestinal stem cells. Following 8 Gy RT, OLFM4-positive cells were significantly reduced by both conventional and FLASH RT (Fig. S6). However, in the control and HI 24-hour groups, FLASH RT resulted in significantly less stem cell loss compared to conventional RT. In contrast, in the HI 96-hour group, the reduction in OLFM4-positive cells was comparable between conventional and FLASH RT. Although stem cell sensitivity to RT was more pronounced after 10 Gy, the high iron diet appeared to diminish the protective effect of FLASH RT. Taken together, our findings suggest that iron availability plays a crucial role in RT-induced lipid peroxidation and contributes to the reduced normal tissue damage observed with FLASH RT.

## Discussion

Approximately 50% of cancer patients undergo RT during their cancer treatment. Advances in RT techniques have resulted in more precise and focused treatments (50, 51). However, the effectiveness of RT is often limited by the radioresistance of tumors and normal tissue toxicities, particularly for tumors located in sensitive areas such as the abdomen or bone marrow. FLASH RT, introduced in 2014 by Fauvadon *et al*. shows promise in enhancing the therapeutic window of RT by decreasing normal tissue toxicity while maintaining tumor control (52). Studies by multiple research groups have further validated FLASH effects using various *in vitro* and *in vivo* models (53). Moreover, the successful application of FLASH dose rates to cancer patients in several clinical trials is promising for its translation into the clinic (54–57).

The most widely proposed mechanism for decreased FLASH RT normal tissue toxicity is oxygen depletion (11, 14–16, 53). Extensive *in vitro* and *in vivo* studies have shown that the sparing effect seen at ultra-high dose rates varies with variable oxygen, supporting the crucial role of oxygen in response to FLASH RT (15–18, 58). However, some of these studies also indicated that FLASH-induced hypoxia alone was not sufficient to achieve normal tissue sparing (17, 18). To fully account for decreased normal tissue toxicity, multiple studies have proposed other potential FLASH effects, including radical-radical recombination, decreased DNA damage, conservation of the stem cell niche, and immune responses (16, 19, 21, 47, 48, 59–62). While these studies suggest that the FLASH effect may arise from the complex interactions of chemical and biological responses within the tissue microenvironment, the underlying cause of the differential effects on tumor and normal tissues remains incompletely elucidated.

In this study, we demonstrate that FLASH RT induced lipid peroxidation and ferroptosis, an iron dependent regulated cell death pathway, based on our *in vitro* clonogenic survival data. Furthermore, we showed that these effects are attributed to differences in iron levels in normal and tumor tissues. Supporting our hypothesis, a previous study by Froidevaux *et al*. showed that FLASH RT did not induce lipid peroxidation, an essential step for ferroptosis, compared to conventional RT, using micelles and liposomes (22). However, this study did not differentiate FLASH effects between tumor vs normal tissues, and did not account for the complexity of the tissue microenvironment. In the present study, we compared the effects of FLASH RT on lipid peroxidation *in vitro* and *in vivo* utilizing several cancer cells, xenograft tumors, and normal tissues. In cancer cells and tumors, we found increased lipid peroxidation compared to non-irradiated controls after conventional and FLASH RT. Moreover, this increase in lipid peroxidation corresponded to an increase in ferroptosis *in vitro*. In contrast, we observed significantly reduced lipid peroxidation in normal tissues after FLASH RT compared to conventional RT, in line with the previous observations regarding FLASH effects on lipid peroxidation (22). Additionally, we observed that elevated lipid peroxidation after conventional RT led to tissue damage via ferroptosis.

Interestingly, we also demonstrated that a variety of RT types (including protons and alpha particles) are capable of inducing lipid peroxidation in cancer cells despite qualitative and quantitative differences in DNA damage they cause. Since the FLASH effect has been seen in studies using proton beams as well as heavier ion beams (helium and carbon beams), our data further support the role of lipid peroxidation for the FLASH effect (18, 48, 60, 63–69).

Iron is essential in cell survival and biogenesis, as well as in rapidly proliferating tumor cells (2). However, if homeostatic levels of iron become excessive, cells will die due to ROS production through the Fenton reaction. In ferroptosis, iron is crucial as it initiates cell death by facilitating ROS production, leading to lipid peroxidation (6). The present study evidenced the crucial role of iron in cancer cells and their dependency on iron transport for survival. We also confirmed that excess iron increased the radiosensitization of tumor cells through ferroptosis, while iron chelation protected against RT-induced ferroptosis. Given our observation of significantly elevated iron levels in lung and breast cancer tumors compared to normal tissues, we hypothesized that baseline tissue iron levels may critically influence the reduction of normal tissue toxicity following FLASH radiotherapy.

To test our hypothesis regarding the role of tissue iron levels in the FLASH effect, we selectively increased iron levels in the intestines without altering systemic iron homeostasis. In the control diet group, the normal tissue protection by FLASH RT was evidenced by decreased lipid peroxidation and more regenerating crypts compared to conventional RT. Interestingly, elevating iron levels only at the time of RT was not sufficient to reverse the FLASH effect, since rapid iron turnover immediately decreased normal tissue iron levels after withdrawal of the high-iron diet. However, when high iron levels were sustained for another 72 hours after IR, the FLASH effect was abolished in the upper intestines, indicating that sufficient iron availability during and after RT is necessary to induce lipid peroxidation leading to tissue damage, which reflects underlying biological processes.

The kinetic model proposed by Spitz *et al*. suggests that the limited Fenton type reaction in normal tissues compared to tumors contributes to the FLASH effect (70). Our study supports this model by demonstrating the variation in intrinsic iron levels between tumors and normal tissues and highlighting the necessity of the prolonged iron presence for the abolishment of the FLASH sparing effect. Future studies exploring the benefits or disadvantages of FLASH effects on iron-rich tissues (such as the liver or bone marrow), focusing on lipid peroxidation and ferroptosis, will be required. In conclusion, the present study highlights that iron, lipid peroxidation and ferroptosis contribute to the FLASH effect, furthering our understanding of the mechanisms underlying reduced toxicity in normal tissue after FLASH RT. Moreover, our study could also be used as a rationale as to which tissues or organs would benefit from FLASH RT.

## Acknowledgements and funding resources

We acknowledge our use of the gene set enrichment analysis, GSEA software, and Molecular Signature Database (MSigDB) (Subramanian, Tamayo, et al. (2005), PNAS 102, 15545-15550, http://www.broad.mit.edu/gsea/). Fig. 5A was created in https://BioRender.com. This work was supported by Medical Research Council (MRC) Program grant (MC_UU_00001/11), Ministry of Health and Welfare in Korea (RS-2023-00266627), National Institute of Health PO1 grant (5P01CA257904-03), Gray Trust (AN9050), Cancer Research UK—RadNet (C6078/A28736), MRC (MR/X006611/1), PhD students Study Abroad Program, National Science and Technology Council (NSTC), Taiwan (114-2917-I-038 -001), and John Fell Fund (AND00180).

## Author contributions

Study Conception & Design: N.V.L., M.H., C.C., S.L., K.P., and E.J.M.; Performed Experiment and Data Collection: N.V.L., J.K, G.N., I.D.C.T., S.P., J.R, P.J.L., Y.J., S.P., T.H., A.N., E.C., E.W., A.A, M.H., C.C., C.K.T., K.P., and E.J.M.; Performed Data Analysis: N.V.L., J.K, G.N., P.J.L., S.P., T.H., A.N., E.C., E.W., M.H., C.C., and E.J.M.; Interpretation of data analysis: N.V.L., J.K, M.H., C.C., S.C.S., I.J.L., Y.T.J., A.J.G., K.P., and E.J.M. ;Writing the first draft: N.V.L. and E.J.M.; Figures Design: N.V.L., J.K., and E.J.M. Supervision: A.J.G., K.P., and E.J.M; Funding acquisition: I.J.L., S.C.S., A.J.G., K.P., and E.J.M. All authors contributed to manuscript editing and revision

## References

1. Federico G, Carrillo F, Dapporto F, Chiariello M, Santoro M, Bellelli R, et al. NCOA4 links iron bioavailability to DNA metabolism. Cell Rep. 2022;40(7):111207.

2. Torti SV, Manz DH, Paul BT, Blanchette-Farra N, Torti FM. Iron and Cancer. Annu Rev Nutr. 2018;38:97–125.

3. Hann HW, Stahlhut MW, Menduke H. Iron enhances tumor growth. Observation on spontaneous mammary tumors in mice. Cancer. 1991;68(11):2407–10.

4. Radulescu S, Brookes MJ, Salgueiro P, Ridgway RA, McGhee E, Anderson K, et al. Luminal iron levels govern intestinal tumorigenesis after Apc loss in vivo. Cell Rep. 2012;2(2):270–82.

5. Sukiennicki GM, Marciniak W, Muszynska M, Baszuk P, Gupta S, Bialkowska K, et al. Iron levels, genes involved in iron metabolism and antioxidative processes and lung cancer incidence. PLoS One. 2019;14(1):e0208610.

6. Dixon SJ, Lemberg KM, Lamprecht MR, Skouta R, Zaitsev EM, Gleason CE, et al. Ferroptosis: an iron-dependent form of nonapoptotic cell death. Cell. 2012;149(5):1060–72.

7. Lei G, Zhang Y, Koppula P, Liu X, Zhang J, Lin SH, et al. The role of ferroptosis in ionizing radiation-induced cell death and tumor suppression. Cell Res. 2020;30(2):146–62.

8. Lang X, Green MD, Wang W, Yu J, Choi JE, Jiang L, et al. Radiotherapy and Immunotherapy Promote Tumoral Lipid Oxidation and Ferroptosis via Synergistic Repression of SLC7A11. Cancer Discov. 2019;9(12):1673–85.

9. Ye LF, Chaudhary KR, Zandkarimi F, Harken AD, Kinslow CJ, Upadhyayula PS, et al. Radiation-Induced Lipid Peroxidation Triggers Ferroptosis and Synergizes with Ferroptosis Inducers. ACS Chem Biol. 2020;15(2):469–84.

10. E. Hall AG. Radiation Biology for Radiologist. 8th Edition ed: Wolters Kluwer Health; 2018.

11. Wilson JD, Hammond EM, Higgins GS, Petersson K. Ultra-High Dose Rate (FLASH) Radiotherapy: Silver Bullet or Fool’s Gold? Front Oncol. 2019;9:1563.

12. Vozenin MC, Bourhis J, Durante M. Towards clinical translation of FLASH radiotherapy. Nat Rev Clin Oncol. 2022;19(12):791–803.

13. Kim MM, Darafsheh A, Schuemann J, Dokic I, Lundh O, Zhao T, et al. Development of Ultra-High Dose-Rate (FLASH) Particle Therapy. IEEE Trans Radiat Plasma Med Sci. 2022;6(3):252–62.

14. Moon EJ, Petersson K, Olcina MM. The importance of hypoxia in radiotherapy for the immune response, metastatic potential and FLASH-RT. Int J Radiat Biol. 2022;98(3):439–51.

15. Adrian G, Konradsson E, Lempart M, Back S, Ceberg C, Petersson K. The FLASH effect depends on oxygen concentration. Br J Radiol. 2020;93(1106):20190702.

16. Montay-Gruel P, Acharya MM, Petersson K, Alikhani L, Yakkala C, Allen BD, et al. Long-term neurocognitive benefits of FLASH radiotherapy driven by reduced reactive oxygen species. Proc Natl Acad Sci U S A. 2019;116(22):10943–51.

17. Cao X, Zhang R, Esipova TV, Allu SR, Ashraf R, Rahman M, et al. Quantification of Oxygen Depletion During FLASH Irradiation In Vitro and In Vivo. Int J Radiat Oncol Biol Phys. 2021.

18. El Khatib M, Van Slyke AL, Velalopoulou A, Kim MM, Shoniyozov K, Allu SR, et al. Ultrafast Tracking of Oxygen Dynamics During Proton FLASH. Int J Radiat Oncol Biol Phys. 2022;113(3):624–34.

19. Jin JY, Gu A, Wang W, Oleinick NL, Machtay M, Spring Kong FM. Ultra-high dose rate effect on circulating immune cells: A potential mechanism for FLASH effect? Radiother Oncol. 2020;149:55–62.

20. Zhu H, Xie D, Wang Y, Huang R, Chen X, Yang Y, et al. Comparison of intratumor and local immune response between MV X-ray FLASH and conventional radiotherapies. Clin Transl Radiat Oncol. 2023;38:138–46.

21. Kim YE, Gwak SH, Hong BJ, Oh JM, Choi HS, Kim MS, et al. Effects of Ultra-high doserate FLASH Irradiation on the Tumor Microenvironment in Lewis Lung Carcinoma: Role of Myosin Light Chain. Int J Radiat Oncol Biol Phys. 2021;109(5):1440–53.

22. Pascal Froidevaux VG, Claude Bailat, Walter Reiner Geyer, François Bochud, Marie-Catherine Vozenin. FLASH irradiation does not induce lipid peroxidation in lipids micelles and liposomes. Radiation Physics and Chemistry. 2023;205.

23. Vilaplana-Lopera N, Abu-Halawa A, Walker E, Kim J, Moon EJ. Ferroptosis, a key to unravel the enigma of the FLASH effect? Br J Radiol. 2022;95(1140):20220825.

24. Grilj V, Paisley R, Sprengers K, Geyer WR, Bailat C, Bochud F, et al. Average dose rate is the primary determinant of lipid peroxidation in liposome membranes exposed to pulsed electron FLASH beam. Radiation Physics and Chemistry. 2024;222.

25. Moon EJ, Mello SS, Li CG, Chi JT, Thakkar K, Kirkland JG, et al. The HIF target MAFF promotes tumor invasion and metastasis through IL11 and STAT3 signaling. Nat Commun. 2021;12(1):4308.

26. Chen EY, Tan CM, Kou Y, Duan Q, Wang Z, Meirelles GV, et al. Enrichr: interactive and collaborative HTML5 gene list enrichment analysis tool. BMC Bioinformatics. 2013;14:128.

27. Kuleshov MV, Jones MR, Rouillard AD, Fernandez NF, Duan Q, Wang Z, et al. Enrichr: a comprehensive gene set enrichment analysis web server 2016 update. Nucleic Acids Res. 2016;44(W1):W90–7.

28. Xie Z, Bailey A, Kuleshov MV, Clarke DJB, Evangelista JE, Jenkins SL, et al. Gene Set Knowledge Discovery with Enrichr. Curr Protoc. 2021;1(3):e90.

29. Subramanian A, Tamayo P, Mootha VK, Mukherjee S, Ebert BL, Gillette MA, et al. Gene set enrichment analysis: a knowledge-based approach for interpreting genome-wide expression profiles. Proc Natl Acad Sci U S A. 2005;102(43):15545–50.

30. Tracy BL, Stevens DL, Goodhead DT, Hill MA. Variation in RBE for Survival of V79-4 Cells as a Function of Alpha-Particle (Helium Ion) Energy. Radiat Res. 2015;184(1):33–45.

31. Berne A, Petersson K, Tullis IDC, Newman RG, Vojnovic B. Monitoring electron energies during FLASH irradiations. Phys Med Biol. 2021;66(4):045015.

32. Ruan JL, Lee C, Wouters S, Tullis IDC, Verslegers M, Mysara M, et al. Irradiation at Ultra-High (FLASH) Dose Rates Reduces Acute Normal Tissue Toxicity in the Mouse Gastrointestinal System. Int J Radiat Oncol Biol Phys. 2021;111(5):1250–61.

33. Petersson K, Jaccard M, Germond JF, Buchillier T, Bochud F, Bourhis J, et al. High dose-per-pulse electron beam dosimetry - A model to correct for the ion recombination in the Advanced Markus ionization chamber. Med Phys. 2017;44(3):1157–67.

34. Vojnovic B, Tullis IDC, Newman RG, Petersson K. Monitoring beam charge during FLASH irradiations. Frontiers in Physics. 2023;11.

35. Choi C, Lee GH, Son A, Yoo GS, Yu JI, Park HC. Downregulation of Mcl-1 by Panobinostat Potentiates Proton Beam Therapy in Hepatocellular Carcinoma Cells. Cells. 2021;10(3).

36. Franken NA, Rodermond HM, Stap J, Haveman J, van Bree C. Clonogenic assay of cells in vitro. Nat Protoc. 2006;1(5):2315–9.

37. Sondka Z, Bamford S, Cole CG, Ward SA, Dunham I, Forbes SA. The COSMIC Cancer Gene Census: describing genetic dysfunction across all human cancers. Nat Rev Cancer. 2018;18(11):696–705.

38. Huang DW, Sherman BT, Tan Q, Kir J, Liu D, Bryant D, et al. DAVID Bioinformatics Resources: expanded annotation database and novel algorithms to better extract biology from large gene lists. Nucleic Acids Res. 2007;35(Web Server issue):W169-75.

39. Cerami E, Gao J, Dogrusoz U, Gross BE, Sumer SO, Aksoy BA, et al. The cBio cancer genomics portal: an open platform for exploring multidimensional cancer genomics data. Cancer Discov. 2012;2(5):401–4.

40. Chandrashekar DS, Bashel B, Balasubramanya SAH, Creighton CJ, Ponce-Rodriguez I, Chakravarthi B, et al. UALCAN: A Portal for Facilitating Tumor Subgroup Gene Expression and Survival Analyses. Neoplasia. 2017;19(8):649–58.

41. Lakhal-Littleton S, Wolna M, Chung YJ, Christian HC, Heather LC, Brescia M, et al. An essential cell-autonomous role for hepcidin in cardiac iron homeostasis. Elife. 2016;5.

42. Jiang Y, Martin J, Alkadhimi M, Shigemori K, Kinchesh P, Gilchrist S, et al. Olaparib increases the therapeutic index of hemithoracic irradiation compared with hemithoracic irradiation alone in a mouse lung cancer model. Br J Cancer. 2021;124(11):1809–19.

43. Schindelin J, Arganda-Carreras I, Frise E, Kaynig V, Longair M, Pietzsch T, et al. Fiji: an open-source platform for biological-image analysis. Nat Methods. 2012;9(7):676-82.

44. Vozenin MC, Hendry JH, Limoli CL. Biological Benefits of Ultra-high Dose Rate FLASH Radiotherapy: Sleeping Beauty Awoken. Clin Oncol (R Coll Radiol). 2019;31(7):407–15.

45. Adrian G, Konradsson E, Beyer S, Wittrup A, Butterworth KT, McMahon SJ, et al. Cancer Cells Can Exhibit a Sparing FLASH Effect at Low Doses Under Normoxic In Vitro-Conditions. Front Oncol. 2021;11:686142.

46. Anderson NM, Simon MC. The tumor microenvironment. Curr Biol. 2020;30(16):R921–R5.

47. Levy K, Natarajan S, Wang J, Chow S, Eggold JT, Loo PE, et al. Abdominal FLASH irradiation reduces radiation-induced gastrointestinal toxicity for the treatment of ovarian cancer in mice. Sci Rep. 2020;10(1):21600.

48. Kim MM, Verginadis, II, Goia D, Haertter A, Shoniyozov K, Zou W, et al. Comparison of FLASH Proton Entrance and the Spread-Out Bragg Peak Dose Regions in the Sparing of Mouse Intestinal Crypts and in a Pancreatic Tumor Model. Cancers (Basel). 2021;13(16).

49. J. F. Collins SRLF, X. Wang, G. J. Anderson. Mechanisms and Regulation of Intestinal Iron Transport. In: Said HM, editor. Physiology of the Gastrointestinal Tract. 6th edition ed: Academic Press; 2018. p. 1451-83.

50. Bernier J, Hall EJ, Giaccia A. Radiation oncology: a century of achievements. Nat Rev Cancer. 2004;4(9):737–47.

51. Delaney G, Jacob S, Featherstone C, Barton M. The role of radiotherapy in cancer treatment: estimating optimal utilization from a review of evidence-based clinical guidelines. Cancer. 2005;104(6):1129–37.

52. Favaudon V, Caplier L, Monceau V, Pouzoulet F, Sayarath M, Fouillade C, et al. Ultrahigh dose-rate FLASH irradiation increases the differential response between normal and tumor tissue in mice. Sci Transl Med. 2014;6(245):245ra93.

53. Limoli CL, Vozenin MC. Reinventing Radiobiology in the Light of FLASH Radiotherapy. Annu Rev Cancer Biol. 2023;7:1–21.

54. Kinj R, Gaide O, Jeanneret-Sozzi W, Dafni U, Viguet-Carrin S, Sagittario E, et al. Randomized phase II selection trial of FLASH and conventional radiotherapy for patients with localized cutaneous squamous cell carcinoma or basal cell carcinoma: A study protocol. Clin Transl Radiat Oncol. 2024;45:100743.

55. Mascia AE, Daugherty EC, Zhang Y, Lee E, Xiao Z, Sertorio M, et al. Proton FLASH Radiotherapy for the Treatment of Symptomatic Bone Metastases: The FAST-01 Nonrandomized Trial. JAMA Oncol. 2023;9(1):62–9.

56. Bourhis J, Sozzi WJ, Jorge PG, Gaide O, Bailat C, Duclos F, et al. Treatment of a first patient with FLASH-radiotherapy. Radiother Oncol. 2019;139:18–22.

57. Gaide O, Herrera F, Jeanneret Sozzi W, Goncalves Jorge P, Kinj R, Bailat C, et al. Comparison of ultra-high versus conventional dose rate radiotherapy in a patient with cutaneous lymphoma. Radiother Oncol. 2022;174:87–91.

58. Ha B, Liang K, Liu C, Melemenidis S, Manjappa R, Viswanathan V, et al. Real-time optical oximetry during FLASH radiotherapy using a phosphorescent nanoprobe. Radiother Oncol. 2022;176:239–43.

59. Eggold JT, Chow S, Melemenidis S, Wang J, Natarajan S, Loo PE, et al. Abdominopelvic FLASH Irradiation Improves PD-1 Immune Checkpoint Inhibition in Preclinical Models of Ovarian Cancer. Mol Cancer Ther. 2022;21(2):371–81.

60. Velalopoulou A, Karagounis IV, Cramer GM, Kim MM, Skoufos G, Goia D, et al. FLASH Proton Radiotherapy Spares Normal Epithelial and Mesenchymal Tissues While Preserving Sarcoma Response. Cancer Res. 2021;81(18):4808–21.

61. Fouillade C, Curras-Alonso S, Giuranno L, Quelennec E, Heinrich S, Bonnet-Boissinot S, et al. FLASH Irradiation Spares Lung Progenitor Cells and Limits the Incidence of Radio-induced Senescence. Clin Cancer Res. 2020;26(6):1497–506.

62. Labarbe R, Hotoiu L, Barbier J, Favaudon V. A physicochemical model of reaction kinetics supports peroxyl radical recombination as the main determinant of the FLASH effect. Radiother Oncol. 2020;153:303–10.

63. Kristensen L, Poulsen PR, Kanouta E, Rohrer S, Ankjaergaard C, Andersen CE, et al. Spread-out Bragg peak FLASH: quantifying normal tissue toxicity in a murine model. Front Oncol. 2024;14:1427667.

64. Daugherty EC, Zhang Y, Xiao Z, Mascia AE, Sertorio M, Woo J, et al. FLASH radiotherapy for the treatment of symptomatic bone metastases in the thorax (FAST-02): protocol for a prospective study of a novel radiotherapy approach. Radiat Oncol. 2024;19(1):34.

65. Ni H, Reitman ZJ, Zou W, Akhtar MN, Paul R, Huang M, et al. FLASH radiation reprograms lipid metabolism and macrophage immunity and sensitizes medulloblastoma to CAR-T cell therapy. Nat Cancer. 2025.

66. Chowdhury P, Velalopoulou A, Verginadis, II, Morcos G, Loo PE, Kim MM, et al. Proton FLASH Radiotherapy Ameliorates Radiation-induced Salivary Gland Dysfunction and Oral Mucositis and Increases Survival in a Mouse Model of Head and Neck Cancer. Mol Cancer Ther. 2024;23(6):877–89.

67. Verginadis, II, Velalopoulou A, Kim MM, Kim K, Paraskevaidis I, Bell B, et al. FLASH proton reirradiation, with or without hypofractionation, reduces chronic toxicity in the normal murine intestine, skin, and bone. Radiother Oncol. 2025;205:110744.

68. Tinganelli W, Weber U, Puspitasari A, Simoniello P, Abdollahi A, Oppermann J, et al. FLASH with carbon ions: Tumor control, normal tissue sparing, and distal metastasis in a mouse osteosarcoma model. Radiother Oncol. 2022;175:185–90.

69. Dokic I, Moustafa M, Tessonnier T, Meister S, Ciamarone F, Akbarpour M, et al. Ultra-High Dose Rate Helium Ion Beams: Minimizing Brain Tissue Damage while Preserving Tumor Control. Mol Cancer Ther. 2024.

70. Spitz DR, Buettner GR, Petronek MS, St-Aubin JJ, Flynn RT, Waldron TJ, et al. An integrated physico-chemical approach for explaining the differential impact of FLASH versus conventional dose rate irradiation on cancer and normal tissue responses. Radiother Oncol. 2019;139:23–7.

