## Supplementary figures for "Tissue-Specific Iron Levels Modulate Lipid Peroxidation and the FLASH Radiotherapy Effect"

Fig. S1

(A)

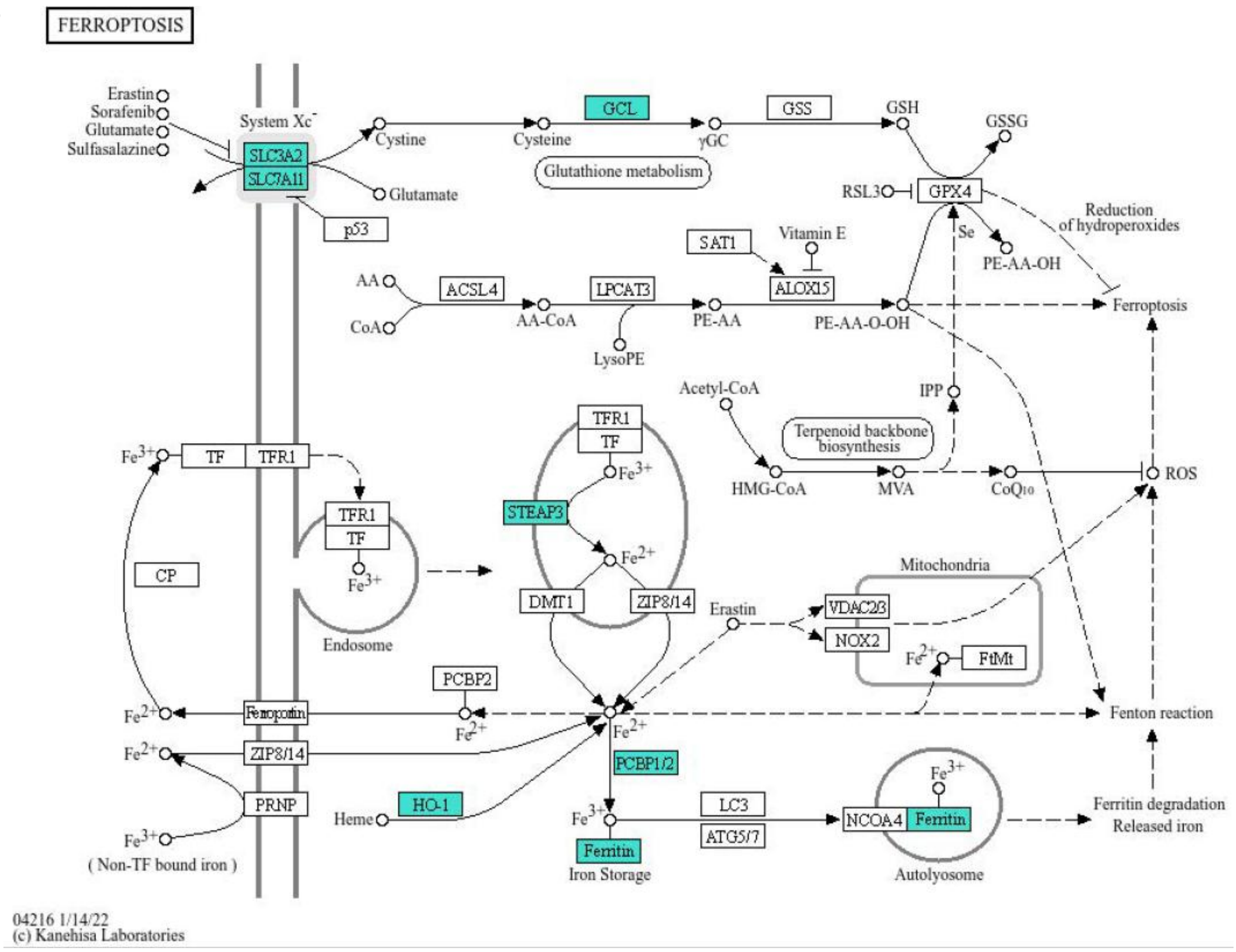


(B)

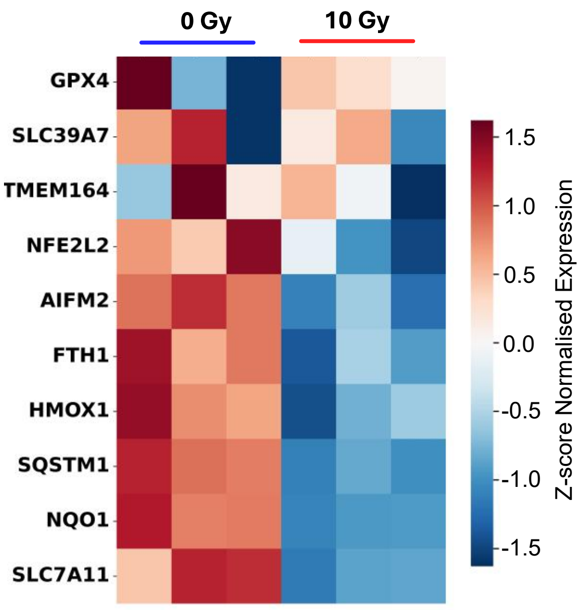


**Fig. S1. Genes in the ferroptosis pathway were significantly altered by radiation.** (A) The list of genes from RNA sequencing was analyzed by Enrichr and those involved in ferroptosis were visualized on the KEGG pathway map. (B) The heatmap generated by GSEA showed a downregulation of genes involved in ferroptosis suppression following radiation.

Fig. S2

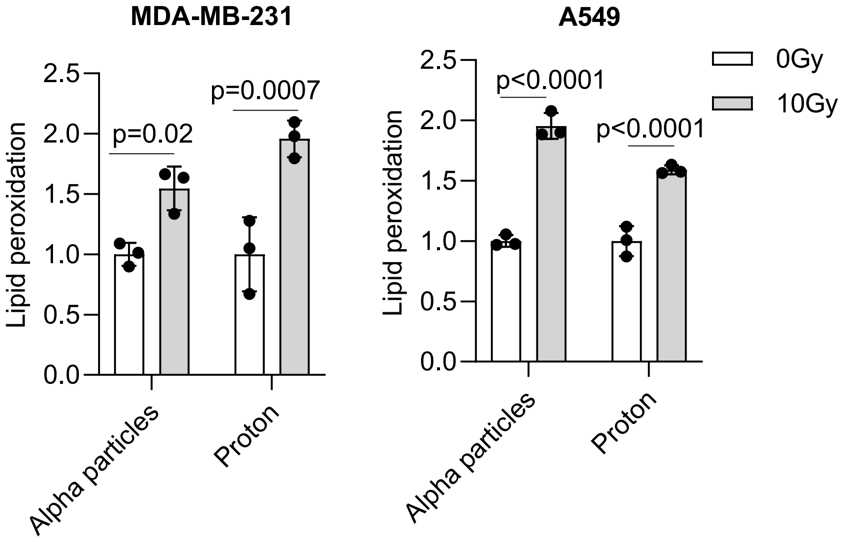


**Fig. S2. Different radiation types similarly induced lipid peroxidation.** MDA-MB-231 and A549 cells were treated with 10 Gy radiation by using alpha-particles or proton. Lipid peroxidation, which was measured 24 hours after RT, was significantly increased by both radiation types (n = 3). Error bars indicate standard deviation (SD). Statistical test was performed by Two-way ANOVA with Šídák's multiple comparisons test.

Fig. S3

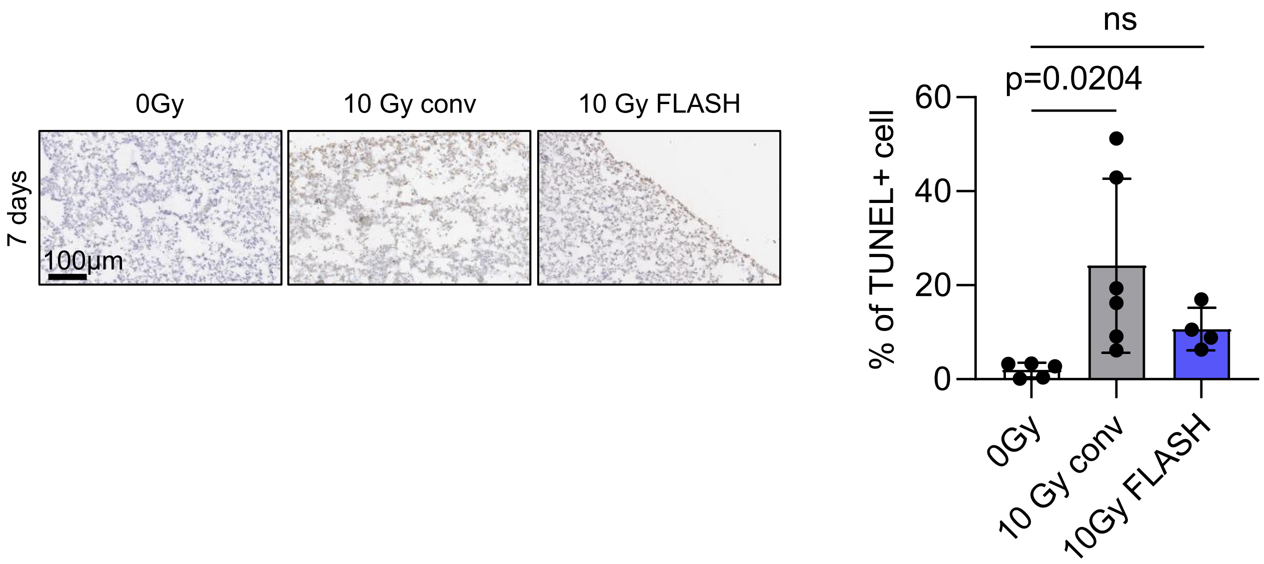


**Fig. S3. Differential apoptosis by conventional and FLASH RT**. Apoptosis in mouse lung tissues was assessed by TUNEL assay 7 days after treatment. While 10 Gy conventional RT significantly increased the population of TUNEL-positive cells, FLASH RT did not induce a comparable level of apoptotic damage.

Fig. S4

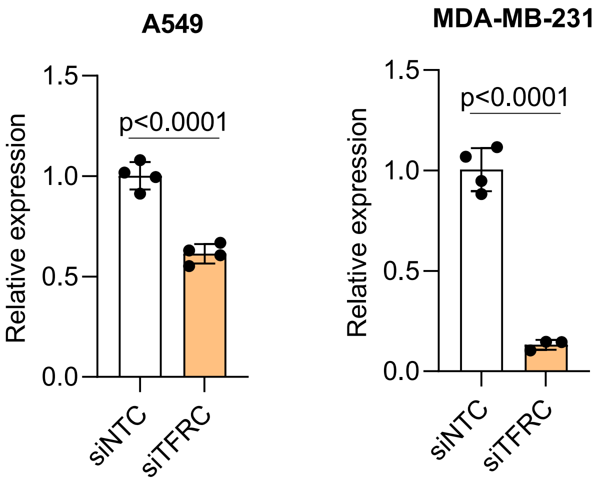


**Fig. S4. Inhibition of TFRC expression**. TFRC expression was inhibited by siRNA transfection in A549 and MDA-MB-231 cells. Significantly reduced expression after knock down was confirmed by qRT-PCR. (n = 3-4, student's *t*-test).

Fig. S5

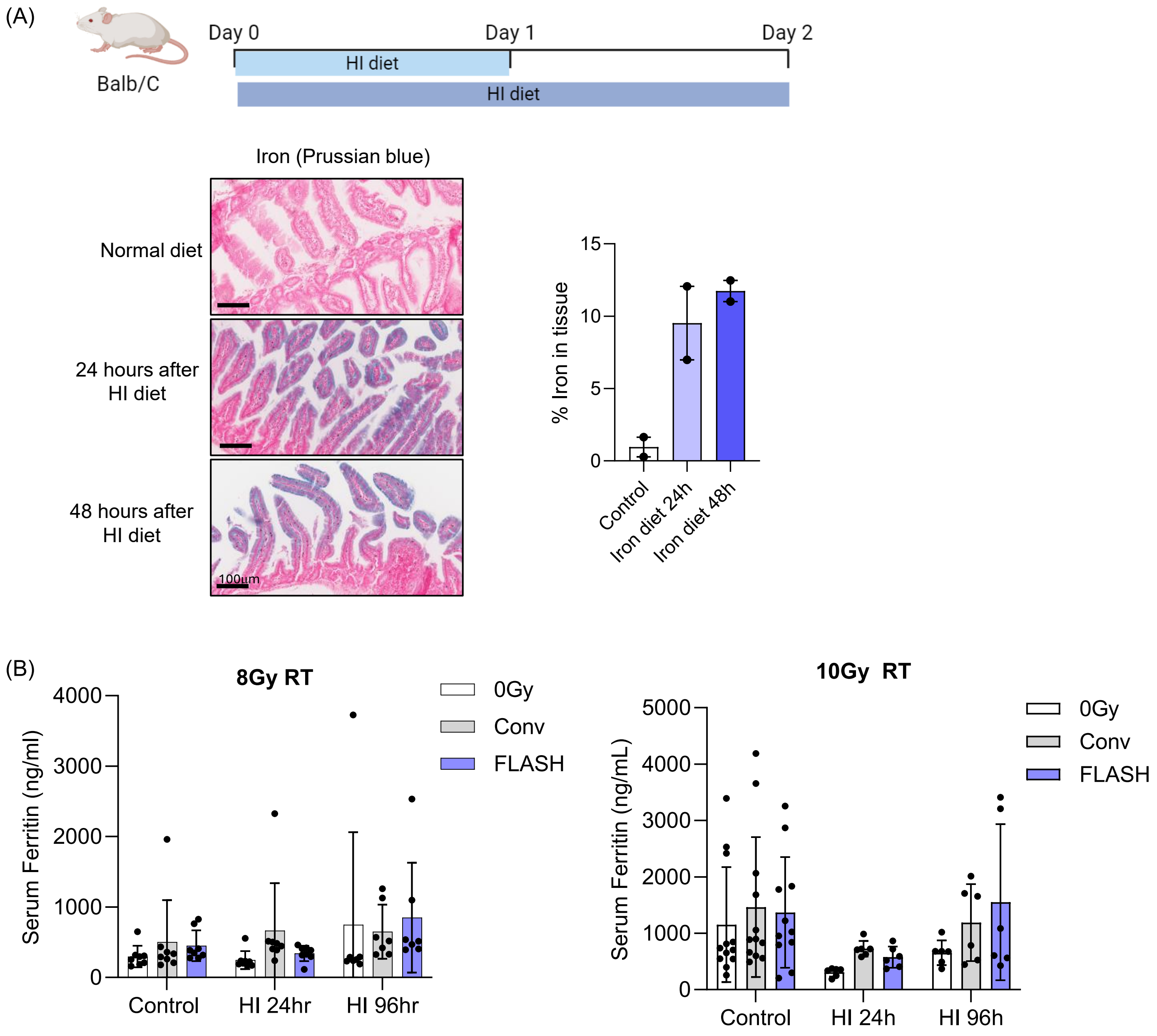


**Fig. S5. FLASH spares intestine tissues and iron levels can be increased in the upper intestines by dietary supplementation.** (A) Increasing dietary iron using an HI diet led to a rapid accumulation of iron in the upper intestines after 24 hours or 48 hours, as measured by using Prussian blue (n = 2). (B) Ferritin levels in the blood were not altered by a high-iron diet, indicating that it does not affect iron levels systemically (For 8 Gy, control diet; n = 8, HI 24-hourl n = 8, HI 96-hour; n=7, For 10 Gy, control diet; n = 12, HI 24-hour; n = 6, HI 96-hour; n = 6). Error bars indicate standard deviation (SD). Statistical tests were performed by One-way ANOVA with Tukey’s multiple comparisons test (B).

Fig. S6

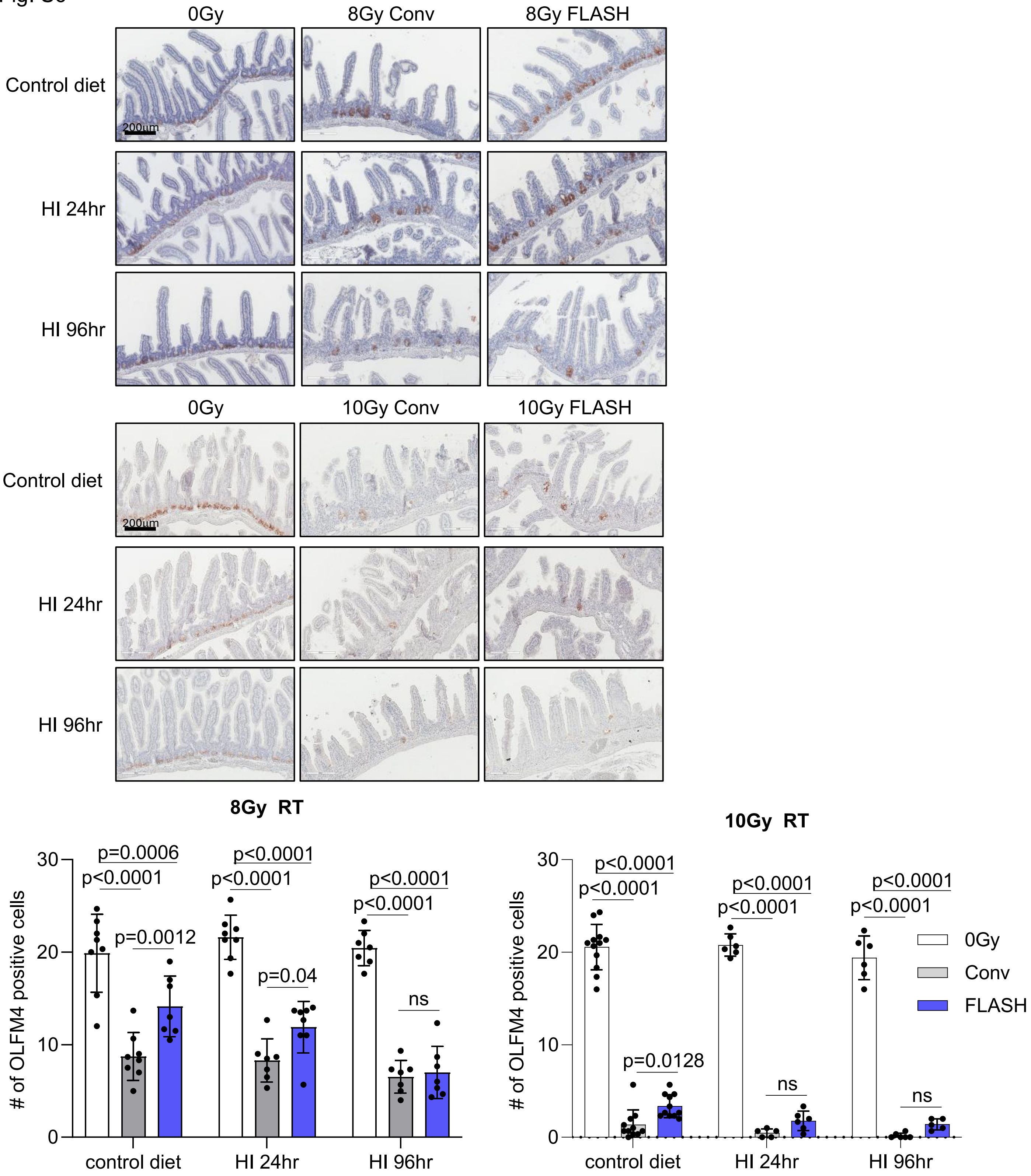


**Fig. S6. Effect of iron diet on intestinal stem cell population.** OLFM4 staining revealed that intestinal stem cells were significantly reduced following both 8 Gy and 10 Gy conventional RT. In contrast, FLASH RT resulted in less reduction of OLFM4-positive cells in the control diet group, indicating a protective effect. However, this protective effect was markedly diminished in mice on a high-iron diet.

Fig. S7

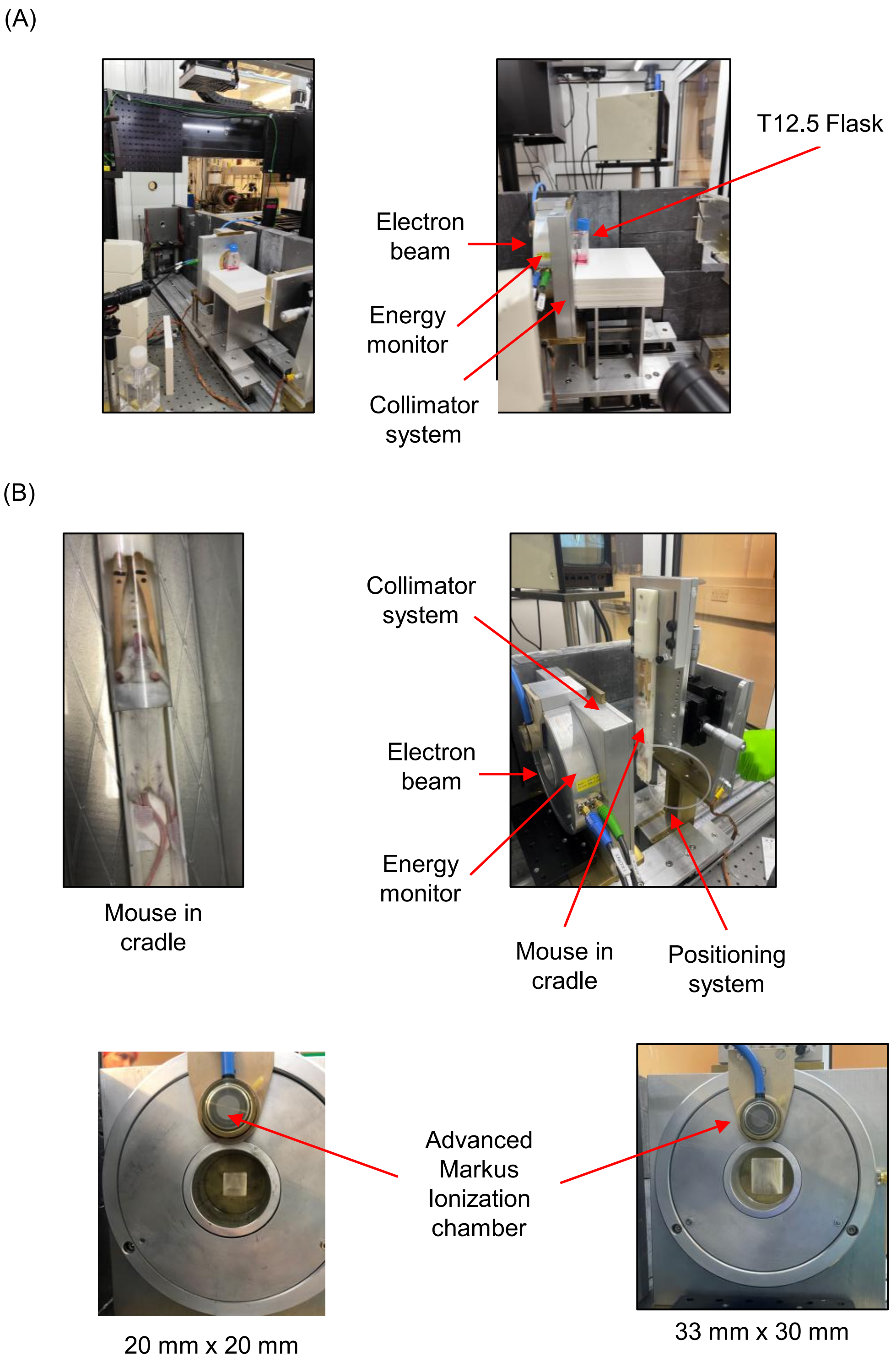


**Fig. S7. FLASH RT is set up to treat both *in vitro* and *in vivo* models.** (A) To perform *in vitro* studies T12.5 flasks or 35 mm cell culture dishes were irradiated in a vertical position in the central part of the beam. (B) The anesthetized mice were placed in an immobilization cradle and irradiated in an upright position. A 6 mm brass collimator was used to define the radiation field covering the whole abdominal area.

Fig. S8

Non-treated

Conventional

FLASH

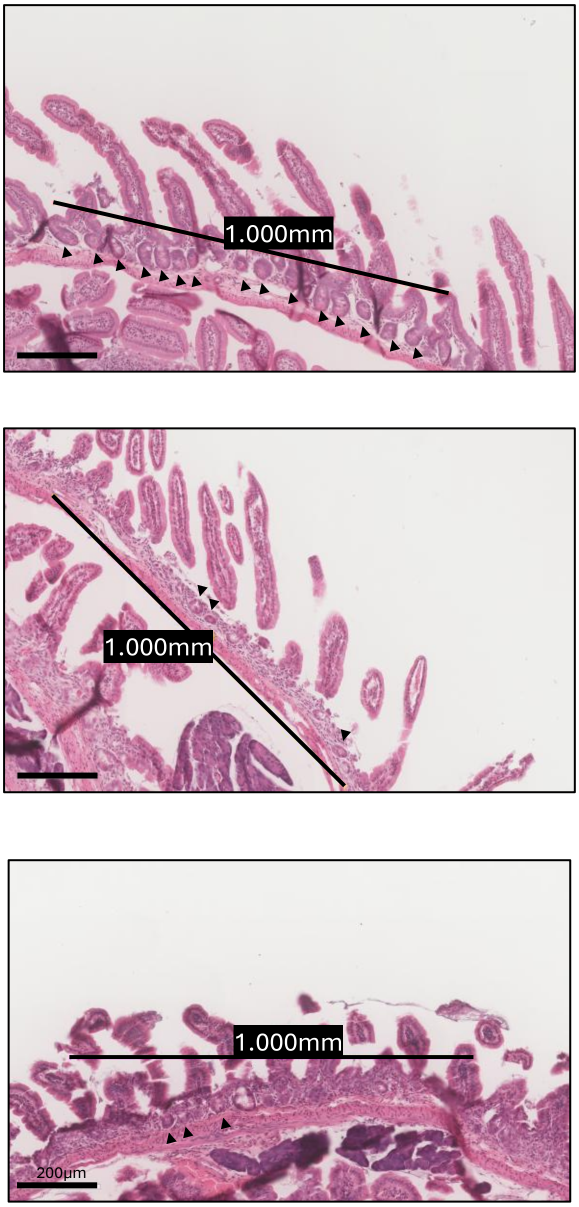


**Fig. S8. The number of remaining crypts indicates radiation-induced damage.** The number of remaining crypts in the upper intestines was quantified by selecting three representative fields, each 1 mm in size, per slide.
